# LAML-Pro: Joint Maximum Likelihood Inference of Cell Genotypes and Cell Lineage Trees

**DOI:** 10.64898/2026.03.27.714879

**Authors:** Gillian Chu, Henri Schmidt, Benjamin J. Raphael

## Abstract

**Motivation:** Recent dynamic lineage tracing technologies use genome editing to induce heritable mutations, or edits, that accumulate across successive cell divisions. These edits are measured using single-cell sequencing or imaging, providing data to reconstruct cell lineages at single-cell resolution. Current computational approaches to infer cell lineage trees, or phylogenies, from these data perform two separate steps: (1) Identify each cell’s edits (genotype) from the raw sequencing or imaging data; (2) Infer a cell lineage tree from the cell genotypes. However, genotyping cells is an inexact process and genotype errors can yield an inaccurate lineage tree. For example, using fluorescence based-imaging to measure edits results in a high fraction (≈ 25-50%) of uncertain or erroneous genotypes.

**Results:** We introduce Lineage Analysis via Maximum Likelihood with PRobabilistic Observations (LAML-Pro), an algorithm that jointly infers cell genotypes and a cell lineage tree. LAML-Pro is based on the Probabilistic Mixed-type Missing Observation (PMMO) model, which we derive to describe both the genome editing and genotype observation processes. LAML-Pro constructs lineage trees from thousands of cells in under an hour by leveraging the sparsity of transitions under the PMMO model. On simulated data, we demonstrate that LAML-Pro corrects genotype errors and infers substantially more accurate trees than existing methods which are vulnerable to genotype errors. Applied to data from two recent imaging-based lineage tracing systems, LAML-Pro reduces genotype errors by 5-fold and produces more spatially coherent lineage trees compared to existing methods.

**Availability and Implementation:** LAML-Pro is freely available at: github.com/raphael-group/LAML-Pro.

## 1 Introduction

Reconstructing the history of cell divisions during the development of an organism is a key goal in developmental biology but remains a major experimental and computational challenge. The history of cell divisions can be summarized in a *cell lineage tree* whose nodes correspond to cells and edges correspond to cell division events. While naturally occurring somatic mutations are heritable markers of cell division and can in principle be used to reconstruct the cell lineage tree, these mutations are difficult and expensive to measure at the necessary coverage and resolution [Lodato et al., 2015]. Thus, dynamic lineage tracing (DLT) systems have been designed to create heritable and non-modifiable edits in the genome as cells divide [Wagner et al., 2020]. In the past few years, multiple dynamic lineage tracing technologies have been developed, improving both the editing process and introducing various approaches to measure the resulting edits [McKenna et al., 2016, Raj et al., 2018, Spanjaard et al., 2018, Alemany et al., 2018, Yang et al., 2022, Chan et al., 2019, Simeonov et al., 2021, Bowling et al., 2020, Tang et al., 2018, Hwang et al., 2019, Chen et al., 2021, Loveless et al., 2021, Choi et al., 2022, Liao et al., 2024, Chow et al., 2021]. Several biological mechanisms are used to create edits for lineage tracing: CRISPR/Cas9-based cut and repair [McKenna et al., 2016, Raj et al., 2018, Spanjaard et al., 2018, Alemany et al., 2018, Yang et al., 2022, Chan et al., 2019, Simeonov et al., 2021, Bowling et al., 2020], CRISPR base editor substitutions [Tang et al., 2018, Hwang et al., 2019], and prime editor-based nick and repair [Chen et al., 2021, Loveless et al., 2021, Choi et al., 2022, Liao et al., 2024]. After edits accumulate in a population of cells, the edits are measured either by DNA sequencing [Masuyama et al., 2019], single cell RNA sequencing [Raj et al., 2018], or fluorescence imaging [Frieda et al., 2017] where the edits fluoresce as different colors and are measured as pixel intensities.

Computational reconstruction of the cell lineage tree from DLT data has been the focus of much recent work and several specialized algorithms have been designed to take advantage of the unique properties of DLT data [Zafar et al., 2020, Jones et al., 2020, Wang et al., 2021, Feng et al., 2021, Sashittal et al., 2022, Seidel et al., 2022, Mai et al., 2024]. Drawing on techniques from species phylogenetics, such specialized algorithms apply distance-based [Wang et al., 2021], maximum parsimony [Jones et al., 2020, Sashittal et al., 2022], and maximum likelihood [Feng et al., 2021, Zafar et al., 2020, Seidel et al., 2022, Mai et al., 2024] techniques to cell lineage tree reconstruction. All current approaches, however, assume that the cell genotypes are provided as input. In fact, cell genotypes are not known and must be inferred prior to running existing cell lineage tree reconstruction algorithms (e.g. using the pipeline in [Jones et al., 2020] to transform observations into edit states or “genotypes”), which is a highly error-prone process [Jones et al., 2024, Koblan et al., 2025].

We introduce Lineage Analysis via Maximum Likelihood with PRobabilistic Observations (LAML-Pro), an algorithm to build cell lineage trees directly from DLT observations (e.g. imaging phenotypes or sequencing reads) by marginalizing over *unknown* cell genotypes. LAML-Pro jointly infers a maximum likelihood tree, model parameters and the cell genotypes under the Probabilistic Mixed-type Missing with Observations (PMMO) model, which integrates a model of the genome editing process with a generative model of the observed data. The approach of jointly inferring the tree and the genotypes from DNA sequencing observations has been demonstrated to improve trees in cancer phylogenetics [Satas et al., 2020, Singer et al., 2018, Kozlov et al., 2022, Chen et al., 2022]; however these cancer phylogenetics methods do not model imaging-based DLT technologies.

We demonstrate LAML-Pro’s advantages on simulated DLT data and two recent imaging-based DLT datasets with high missing data rates of 22–37.5% and uncertainty in the genotype observations (2–35% “low confidence” genotypes [Chadly et al., 2024, Koblan et al., 2025]). LAML-Pro corrects genotype errors and reconstructs more accurate cell lineage trees than existing approaches on DLT data. LAML-Pro reduced genotype errors from imaging-based readout (≈ 25–50%) to the level of *sequencing error* (≤0.03% [Wall et al., 2014]), broadly improving the feasibility of imaging-based readout for DLT systems. Compared to current probabilistic approaches [Chu et al., 2025, Seidel et al., 2022], LAML-Pro scales to data with thousands of cells; requiring e.g. ≈ 2.6 seconds per NNI iteration on a tree with 3,108 cells.

## 2 Methods

A dynamic lineage tracing experiment starts with a single cell containing *K* genomic sites in an unedited state. As the cell and its progeny divide, sites in each cell are edited, resulting in a cell lineage tree 𝒯 = (*V, E*) and *genotypes Z*(*u*) representing the states at the *K* genomic sites in each cell *u* ∈ *V* . Importantly, in most lineage-tracing systems, the edits are non-modifiable, meaning that once a site is edited in a cell, the site cannot be edited again in the descendants of this cell [Sashittal et al., 2022]. At the end of the lineage tracing experiment, the (hidden) genotypes *Z*(*u*) at the extant cells (i.e. leaves in 𝒯) are imperfectly measured, emitting observations *X*(*u*). In particular, the true genotypes *Z*(*u*) at the leaves of the tree 𝒯 are *unknown*, which is a departure from existing models [Chu et al., 2025, Seidel et al., 2022, Sashittal et al., 2022, Jones et al., 2020] of lineage tracing data.

We introduce the Probabilistic Mixed-type Missing with Observations (PMMO) model to describe both the editing process, the different types of missing data, and the relationship between unknown genotypes *Z* and their observations *X*. We also derive an algorithm, Lineage Analysis via Maximum Likelihood with PRobabilistic Observations (LAML-Pro), to simultaneously infer the cell lineage tree 𝒯 and marginalize over the unknown genotypes *Z*. The LAML-Pro algorithm reconstructs the maximum likelihood cell lineage tree 𝒯 = (*V, E*) and PMMO parameters Θ by marginalizing the unknown genotypes of the extant cells (Fig. 1). The edges of 𝒯 represent cell divisions and {*δ*_*e*_}_*e*∈*E*_ are branch lengths representing either elapsed time (i.e. real time between cell divisions) or mutation units (i.e. expected number of mutations between cell divisions). We assume as a prerequisite that 𝒯 is a rooted tree with root vertex *r*_𝒯_ and leaf set *V*_*L*_. The root *r*_𝒯_ of 𝒯 has exactly one child; all other internal (non-leaf) nodes of 𝒯 have exactly two children. After 𝒯 and Θ are inferred, the unknown genotypes *Z* are estimated as the most probable assignment of genotypes to the leaves of 𝒯.

**Figure 1.**
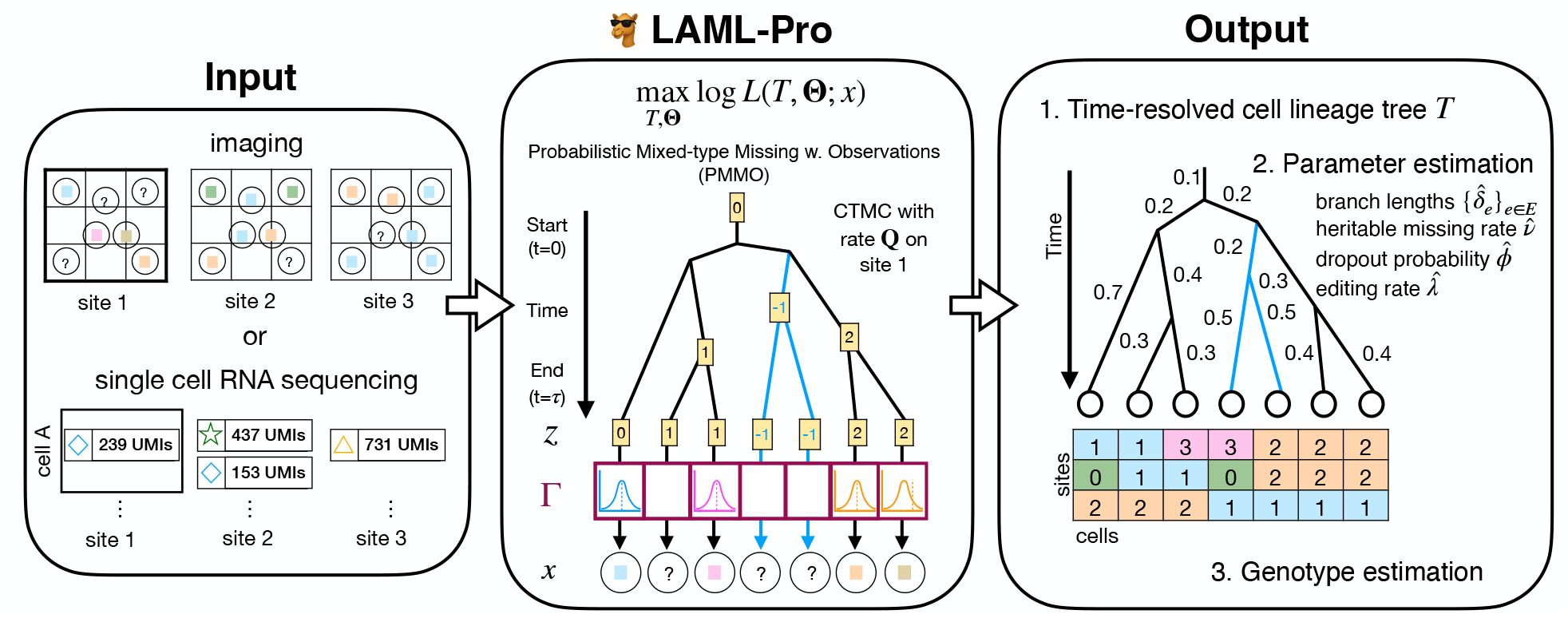
Overview of LAML-Pro. We illustrate two kinds of inputs: fluorescence pixel intensity measurements from imaging or single cell sequenced transcripts with unique molecular identifiers (UMI counts). Given observations (either intensities or UMI counts), LAML-Pro infers the maximum likelihood tree and parameters under the PMMO model. LAML-Pro has three outputs: (1) a maximum likelihood time-resolved cell lineage tree 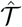; (2) maximum likelihood estimates of other parameters in the PMMO model, including the branch lengths {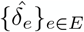}, the heritable missing rate 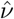, the dropout probability 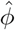, editing rate 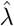; and (3) estimated genotypes.

Formally, LAML-Pro solves the optimization problem

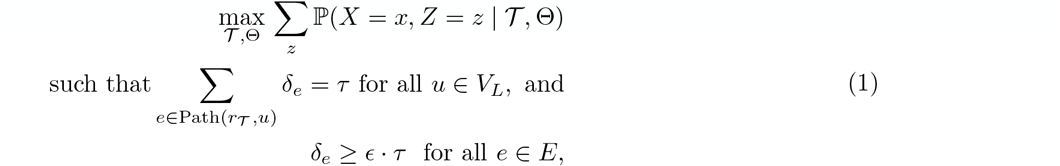

The first constraint in Eq. 1 enforces the strict molecular clock assumption, resulting in an *ultrametric* tree 𝒯 with all root-to-leaf distances equal to *τ* . This constraint is motivated by the observation that in a typical DLT experiment all observed cells are measured at the same time *τ*, and follows previous work in inference of cell lineage trees [Chu et al., 2025, Prillo et al., 2025]. LAML-Pro also solves the optimization problem without the ultrametric constraint, which may be more appropriate when cells are sampled at different times or have heterogeneous division or editing rates.

The second constraint in Eq. 1 imposes a user-specified minimum branch length that scales proportionally with the estimated tree height *τ* (default ∈ = 0.01) as originally proposed in [Prillo et al., 2025]. This constraint serves as a biologically realistic minimum time between cell divisions and was previously shown [Prillo et al., 2025] to improve branch length estimation. Without the branch length lower bound, the true maximum likelihood solution may include zero-length branches when there are no informative edits. However, zero-length branches imply instantaneous cell division, which is both biologically implausible and numerically unstable.

We now describe the PMMO model and the LAML-Pro algorithm. As is standard in statistical phylogenetics, we assume the genotypes evolved independently across the *K* genomic sites [Whelan et al., 2001] and describe the model and likelihood only for a single site.

### 2.1 Probabilistic Mixed-type Missing with Observations (PMMO) Model

#### 2.1.1 Genome editing process

The hidden state/genotype of cell *u* ∈ *V* is denoted by the random variable *Z*(*u*). The hidden alphabet is denoted as {0, 1,… *M*, –1}, where 0 is the *unedited* state, {1, …, *M*} are the possible *edited* states, and –1 is the *heritable missing* state. Biologically, the heritable missing state –1 occurs due to epigenetic silencing or resection [Chan et al., 2019] and results in no observation. As is typical in statistical phylogenetics [Whelan et al., 2001], we model this process with a continuous-time Markov chain (CTMC) that evolves the hidden states along each edge of tree 𝒯. That is, our model of genome editing describes the probability of transitioning between hidden states *Z*(*u*) and *Z*(*v*) along each edge *e* = (*u, v*) ∈ *E*.

Formally, the hidden states evolve according to a CTMC with transition rate matrix

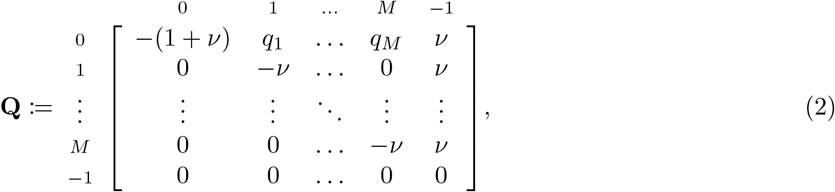

where *q*_1_, *q*_2_, …, *q*_*M*_ are the transition rates from the unedited state 0 to the edited states 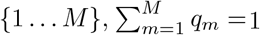, and *v* is the heritable missing rate. In practice, the rates *q*_1_, *q*_2_, …, *q*_*M*_ are either provided as input [Jones et al., 2020, Chu et al., 2025] or assumed to be uniform. The structure of the transition rate matrix in Eq. 2 encodes the non-modifiability and irreversibility of the CRISPR-Cas9 editing process. Then the transition probability from state *a* to *b* along an edge *e* = (*u, v*) with branch length *δ* is given by 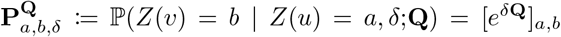, where *e*^**A**^ denotes the matrix exponential of a square matrix **A**. Interestingly, the matrix exponential *e*^*δ***Q**^ has a closed form under the PMMO model [Chu et al., 2025] (see Eq. S1 in the Supplement).

#### 2.1.2 Observation process

The hidden state for an extant cell *u* ∈ *V*_*L*_ is denoted *Z*(*u*). Let the corresponding observation be denoted with the random variable *X*(*u*). Following standard convention, we let *x*(*u*) and *z*(*u*) be instances of the random variables *X*(*u*) and *Z*(*u*), and let *X* be the set of all observations (resp. denote hidden states as *Z*). The observation model asserts that the observations *X* are independent conditioned on the hidden states *Z*, i.e., 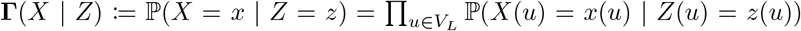. We denote missing observations by the symbol ‘?’, and the dropout probability as *ϕ*. The PMMO model is completely specified by observation probabilities/densities **Γ**(*x* | *z*) := ℙ (*X*(*u*) = *x* | *Z*(*u*) = *z*), where

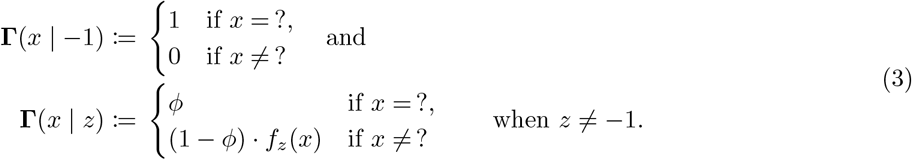

Here *f*_*z*_(*x*) is the probability/density of generating a (non-missing) observation *x* from the (unsilenced) hidden state *z*. Thus, we have presented a general framework for the observation model. Next, we derive the observation model for imaging-based data. See Supp. Sec. S1.2 for the observation model for sequencing-based read count observations and Supp. Sec S1.3 for the character matrix setting.

##### Generating imaging-based observations

Following the definition of the observation model density Γ(*x*|*z*) in Eq. 3, it suffices to specify the conditional density *f*_*z*_(*x*) of generating an imaging observation *x* from the hidden label *z*. To determine *f*_*z*_, we train a kernel density estimator from a set of labeled imaging observation and character state pairs 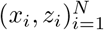 as detailed below.

The imaging observations are represented as points *x*_*i*_ in ℝ^*d*^ where *d* is the number of measured features. The measurements represent *d* statistics derived directly from fluorescence-based readouts [Chadly et al., 2024, Koblan et al., 2025]. To obtain *f*_*z*_, we train *M* + 1 Gaussian kernel density estimators by partitioning the labeled training data 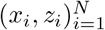 using the hidden state label *z*_*i*_ ∈ {0, 1,…, *M* } and train a separate estimator for each partition.

Labeled imaging observation and character state pairs 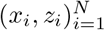 are provided as part of the DLT system design. Crucially, such labeled training data is always provided in imaging-based DLT systems as it is necessary to assign the (hidden) character state *z* to each (apriori unlabeled) imaging observation *x*. We adhere closely to the published classification procedure [Chadly et al., 2024] when training the *M* + 1 kernel density estimators *f*_*z*_. Specifically, we learn a single set of densities for sites that share a common alphabet [Chadly et al., 2024], whereas we learn site-specific sets of densities for site-specific alphabets (consistent with the published use of site-specific classifiers) [Koblan et al., 2025]. See Supp. Secs. S2.4.1 and S2.5.1 for hyperparameter choices (e.g. bandwidth, PCA dimension).

### 2.2 LAML-Pro: Maximum likelihood estimation

In general, maximum likelihood inference of phylogenetic trees is NP-hard [Roch, 2006]. LAML-Pro performs a heuristic maximum likelihood tree search by alternating between (i) proposing a new tree topology and (ii) optimizing model parameters given the proposed topology. Starting from an initial tree (user input, or Neighbor Joining by default), LAML-Pro proposes a candidate topology 𝒯′ using a nearest neighbor interchange (NNI) move [Robinson, 1971, Moore et al., 1973]. Given the proposed topology, LAML-Pro optimizes model parameters 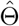 by maximizing the log-likelihood log *L*(𝒯′,Θ;*x*) using an efficient EM algorithm. Proposed trees are accepted or rejected according to a simulated annealing temperature schedule and this process is repeated for a fixed number of iterations. LAML-Pro is summarized in Algorithm 1.

There are two key algorithmic differences compared to the optimization procedure in LAML [Chu et al., 2025]: (i) log-likelihoods are computed under the PMMO observation model and (ii) parameter optimization within the EM algorithm is improved.

#### 2.2.1 Likelihood computation

In the following, we take the estimated parameters as Θ = {*v, φ, τ* }∪ {*δ*_*e*_}_*e*∈*E*_ and all remaining parameters (i.e. *E, q*_1_,…, *q*_*M*_ ) are held fixed. Given the tree topology 𝒯 and parameters Θ, the log-likelihood log *L*(𝒯, Θ; *x*) of the observed data *x* under PMMO is equal to the log of the probability ℙ (*X* = *x* | 𝒯, Θ). As observations at the leaves are conditionally independent given the corresponding hidden states [Felsenstein, 2004], and the hidden states evolve according to the Markov process detailed in PMMO, we marginalize over all assignments of the hidden states *Z* so that ℙ (*X* = *x* |𝒯, Θ) = Σ_*z*_ ℙ (*X* = *x, Z* = *z* |𝒯, Θ). Then,

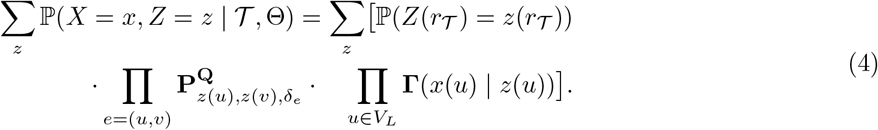

In the preceding, ℙ (*Z*(*r*_𝒯_) = *z*(*r*_𝒯_)) is the probability distribution at the root which is assumed to be fixed with ℙ(*Z*(*r*_𝒯_) = 0) = 1. This likelihood is computed in 𝒪 (*NM*^2^) time by Felsenstein’s pruning algorithm [Felsenstein, 2004], and is further improved to 𝒪 (*NM* ) using the sparsity of the exponentiated transition rate matrix as proposed in [Chu et al., 2025].

#### 2.2.2 EM algorithm for likelihood optimization

We use an expectation maximization (EM) algorithm to optimize the parameters on a given fixed tree topology under the PMMO model. The complexity of the E and M-steps under the PMMO model is the same as in LAML [Chu et al., 2025]. While LAML optimizes parameters sequentially using block coordinate ascent [Chu et al., 2025], we jointly estimate all parameters using an interior point method [Nocedal et al., 2006, Wächter et al., 2006, Biegler et al., 2009]. The advantage of using an interior-point method as opposed to block coordinate ascent [Chu et al., 2025] is that we obtain fast quadratic convergence to stationary (KKT) points, which was not guaranteed by LAML. Further, by making use of the sparsity of the Hessian matrix in the M-step, the cost of using a second-order method as opposed to a first-order method is insubstantial. Finally, we enforce the biologically-motivated minimum branch length constraint in the M-step, which improves both numerical stability and interpretability of inferred trees, as proposed in [Prillo et al., 2025].

Let Θ^*t*^ be the estimated parameters at iteration *t* of the algorithm. At each iteration, the standard EM algorithm maximizes the following lower bound on the log-likelihood:

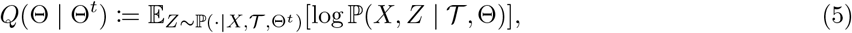

where the expectation is taken with respect to the conditional distribution of hidden variables *Z* given the observed data *X* under current parameter estimates Θ^*t*^. Leveraging previously introduced terms [Mai et al., 2024], we extend the prior derivation of the complete log-likelihood in [Siepel, 2002] to include an observation model on the leaf states, so that *Q*(Θ | Θ^*t*^) has the following form under the PMMO model (up to a constant additive constant)

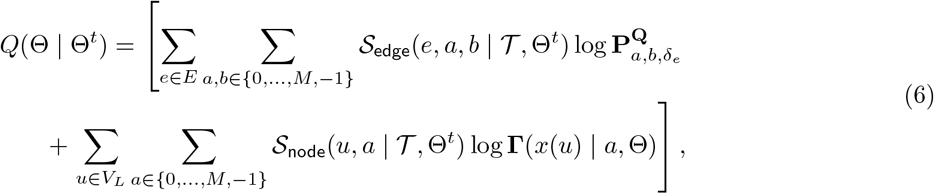

where 𝒮_edge_((*u, v*), *a, b*, | 𝒯, Θ) := ℙ(*Z*(*u*) = *a, Z*(*v*) = *b* | *X* = *x*, 𝒯, Θ) is the posterior transition and 𝒮_node_(*u, z* | *T*, Θ) := ℙ(*Z*(*u*) = *z* | *X* = *x*, 𝒯, Θ) is the posterior emission expectations. Eq. 6 is derived in Supp. Sec. S1.4.

##### E-step

In the E-step, we compute 𝒮_edge_(*e, a, b* | 𝒯, Θ) – the expected number of transitions from hidden state *a* to hidden state *b* across edge *e* conditioned on the observed data *X* – and 𝒮_node_(*u, z* | *T*, Θ) – the probability that leaf *u* takes hidden state *z* conditioned on the observed data. These quantities are computed efficiently via Felsenstein’s pruning algorithm [Felsenstein, 2004] followed by a top-down tree traversal in 𝒪 (*NM* ^2^) time for *N* observed cells and an alphabet of size *M* + 2. Leveraging the sparsity of the exponentiated transition rate matrix, we improve the complexity of the E-step to 𝒪 (*NM* ) as proposed in [Chu et al., 2025].

##### M-step

At iteration *t*, the M-step jointly updates all model parameters via 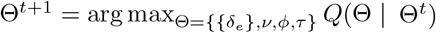 subject to the constraints in Eq. 1. To solve this optimization problem, we provide the interior-point solver IPOPT [Biegler et al., 2009] with analytical gradients and an exact Hessian matrix. Due to the structure of PMMO, the Hessian matrix is sparse and contains only 𝒪 (*N* + *M* ) nonzero elements, where *N* is the number of branches in 𝒯 and *M* + 2 is the alphabet size. All computations are performed in log-space to maintain numerical stability. We provide detailed update equations in Supp. Sec. S1.4.1.

During optimization, branch lengths {*δ*_*e*_}_*e*∈*E*_ are subject to the ultrametric and minimum branch length constraints (Eq. 1). We implicitly estimate the height of the tree *τ* . Since each estimated branch length *δ* is a convolution of the editing rate *λ* and *t* the time, *δ* = *λt* where *δ* is in “edit units” (or so-called “mutation units”). When the experiment length *τ* is given, we compute the editing rate *λ* and rescale the LAML-Pro tree into a chronogram (i.e. branch lengths in time units).

##### Genotype estimation

After inferring a maximum likelihood tree topology 𝒯 and parameters Θ, the maximum a posteriori (MAP) state (equivalently, “genotype”) is 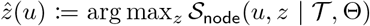, where *z* ∈ {0, 1,…, *M*, ™1}, which corresponds to the most probable hidden state given the observations *x*, tree 𝒯, and parameters Θ. LAML-Pro provides users with MAP genotypes for all nodes in the tree.

## 3 Results

### 3.1 LAML-Pro accurately estimates trees on simulated data

We benchmark LAML-Pro (ultrametric) and LAML-Pro (unconstrained) against four algorithms for single cell lineage tree estimation: LAML on the observed character matrix [Chu et al., 2025], LAML with the ground truth genotypes (True CM), Neighbor Joining (NJ) with the Hamming distance (HD) [Saitou et al., 1987], and NJ with the weighted Hamming distance (wHD) used in [Koblan et al., 2025]. We compare the genotypes estimated by LAML-Pro against the genotypes obtained by calling genotypes prior to tree inference. Finally, we compare LAML-Pro (ultrametric) and LAML-Pro (unconstrained) branch length estimates to six other algorithms: NJ (HD) [Saitou et al., 1987], NJ (wHD) [Jones et al., 2020, Koblan et al., 2025], LAML, LAML (True CM), and ConvexML [Prillo et al., 2025] on NJ-wHD and NJ-HD as in [Koblan et al., 2025].

All methods were evaluated on simulated trees with 12 cell division generations (4,096 cells) using the birth-only process implemented in Cassiopeia [Jones et al., 2020]. We simulated sequence evolution along 400 sites with the alphabet {0, 1, 2, 3, −1} under the PMMO model, matching a realistic lineage tracing dataset [Chadly et al., 2024]. We constructed two model conditions: one without missing data (*v* = 0, *ϕ* = 0) and one with missing data parameters (*v* = 0.15, *ϕ* = 0.1875) shown in Figure 2, calibrated to match 37.5% sites observed as missing data [Chadly et al., 2024]. Observations were drawn from a mixture of 4 spherical Gaussians with means fixed to the vertices of a regular tetrahedron with unit edge length and variance *σ*^2^ (Fig. 2A). We varied the Gaussians’ variance *σ*^2^ to control the genotype error, creating 8 model conditions (Supp. Table S1). Finally, we sampled *n* = 100, 200, 300 cells uniformly at random (without replacement) and took the contracted tree 𝒯′ to be ground truth. Further details are in Supp. Sec. S2.2.

**Figure 2.**
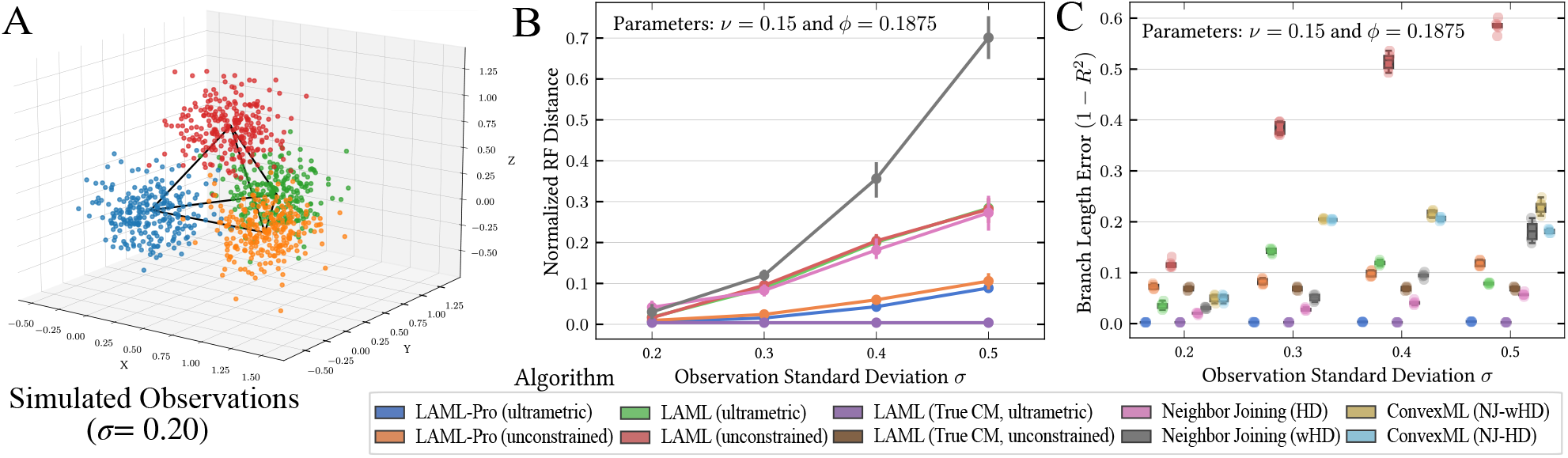
Comparison of LAML-Pro and several other methods on simulated data with 300 cells and ≈ 37.5% missing data (heritable missing rate *v* = 0.15, dropout probability *ϕ* = 0.1875). (A) Observations in three dimensions are drawn from one of four multivariate Gaussians with variance *σ*^2^ centered on the vertices of a regular tetrahedron. (B) Evaluation of methods’ estimated topologies using mean normalized Robinson-Foulds (RF) distance from the true tree, across four model conditions. Error bars indicate standard deviation across five random replicates. (C) Evaluation of error in estimated branch lengths compared to branch lengths on the true tree, across four model conditions. ConvexML infers branch lengths on a given topology.

#### LAML-Pro infers accurate tree topologies and genotypes

LAML-Pro infers more accurate tree topologies than LAML and NJ across all model conditions. With biologically realistic rates of missing data present (37.5% of all cells and sites), both ways of running LAML-Pro substantially reduce the normalized Robinson-Foulds (RF) distance compared to LAML, NJ-HD, and NJ-wHD (median RF LAML-Pro 0.03; LAML 0.18; NJ-HD 0.12; NJ-wHD 0.14; Fig. 2B; Supp. Fig. S2). When no missing data (i.e. *ϕ* = *v* = 0) is present, we observe an even greater improvement (median RF LAML-Pro 0.01; LAML 0.12; NJ-HD 0.07; NJ-wHD 0.08). LAML-Pro (ultrametric) has better performance than LAML-Pro (unconstrained), with lower RF distance (median RF range over model conditions: LAML-Pro (ultrametric) [0.0, 0.178], LAML-Pro (unconstrained) [0.01, 0.263]). LAML-Pro (unconstrained) also outperforms other methods, demonstrating that LAML-Pro is robust to misspecification in the ultrametric assumption. Notably, although NJ (wHD) is widely used [Koblan et al., 2025], NJ (HD) trees incur *much lower* RF error in the presence of heritable missing data. See Supp. S2.2.1 for additional details. LAML-Pro improves over other methods by a greater margin as the observation variance *σ*^2^ increases, due to LAML-Pro’s ability to correct genotype errors (Fig. 2B; Supp. Fig. S3). LAML-Pro achieves similar tree accuracy to a baseline reconstruction method (LAML) that is provided the true genotypes (median RF 0), suggesting LAML-Pro is effective at reconstructing lineage trees despite not directly observing the true cell genotypes.

LAML-Pro correctly infers genotypes and model parameters even as the observation variance *σ*^2^ increases. Specifically, LAML-Pro infers correct genotypes at 90% of sites, while a baseline of picking the most likely genotype (Argmax CM) independently across target sites only infers correct genotypes at 77% of sites.

The improvement of LAML-Pro over this baseline becomes more pronounced as the standard deviation *σ* increases (Supp. Fig. S3). In terms of model parameters, LAML-Pro correctly identifies dropout rate *ϕ* and slightly underestimates heritable missing rate *v* ( Supp. Fig. S4, S5).

LAML-Pro (ultrametric) estimates more accurate branch lengths compared to LAML-Pro (unconstrained), LAML variants, NJ and ConvexML. For example, the branch lengths estimated by LAML-Pro (ultrametric) have nearly perfect (*R*^2^ = 0.995) correlation with the true branch lengths, whereas the branch lengths produced by the next best performing methods NJ-HD and ConvexML (*R*^2^ =0.961, 0.863) are substantially worse (Fig. 2D; Supp. Fig. S5). As the variance *σ*^2^ increases, the branch length estimates by LAML-Pro remain accurate while the branch length estimates degrade for other methods. Notably, even though the branch length estimation method ConvexML [Prillo et al., 2025] is specifically designed for lineage tracing data, it struggles to accurately infer branch lengths in the presence of genotype errors (median *R*^2^ = 0.863).

### 3.2 LAML-Pro corrects genotype errors and improves spatial concordance on PEtracer

We analyze a dataset of 4T1 cells (murine stage IV human breast cancer model) lineage traced with a prime-editing based lineage tracer named PEtracer [Koblan et al., 2025]. Each site has 9 possible states (8 possible edit states and 1 unedited state), and the accumulated edits are read out *in situ* as pixel intensities using FISH imaging across 9 color channels. We focus on three experiments performed using the PEtracer system (Supp. Sec. S2.4). Following the published ultrametric PEtracer trees, we ran LAML-Pro (ultrametric).

#### LAML-Pro corrects imaging-based genotype errors to achieve sequencing-level accuracy

We first evaluate LAML-Pro genotype accuracy on the PEtracer-sequencing experiment, where cell edits are predefined with an adapted PEtracer system to enable paired imaging and sequencing observations at every site [Koblan et al., 2025]. We use the imaging-based observations to genotype the clone with the most sites (153 cells from Clone 2 across 60 sites) and validate these calls against the “ground-truth” sequencing-based genotype calls (subset to the sites that were sequenced correctly). We compare the genotype accuracy of LAML-Pro against four baselines: taking the genotype with the brightest fluorescence (a baseline used in the PEtracer paper), the genotype predicted by the PEtracer discriminative classifier, the genotype predicted by the input generative probabilities (LAML-Pro inputs), and the genotypes output by LAML-Pro. This comparison favors the PEtracer genotype calls, since the PEtracer discriminative classifier is trained on all available benchmark data (i.e. amounts to training error). Despite this, LAML-Pro achieved the lowest genotype error, improving error rates to be comparable to sequencing error (LAML-Pro= 0.3% genotype errors, Fig. 3A), narrowing the gap between sequencing and imaging-based observation accuracy. Visually, LAML-Pro corrects spurious genotype errors (Supp. Fig. S11), and further imputes genotypes at a fraction of the sites where neither sequencing nor imaging yielded observations, correctly genotyping these sites to match the predefined lineage markers (Supp. Fig. S10).

**Figure 3.**
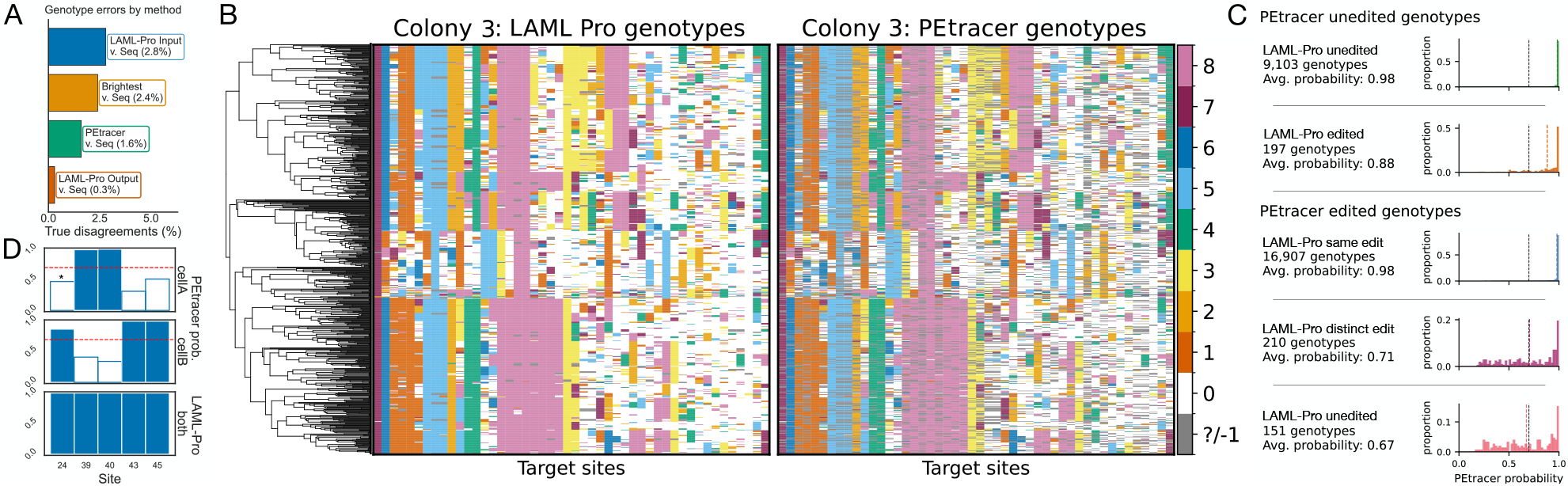
PEtracer. (A) PEtracer-sequencing. Genotype error rates from LAML-Pro and several other methods on imaging-based observations, compared to paired sequencing-based data. (B) PEtracer-colonies. Inferred LAML-Pro tree and genotypes compared to PEtracer genotypes. Cells (rows) in the heatmap match tree taxa and site order is preserved. (C) PEtracer-colonies. Genotype call differences between LAML-Pro and PEtracer, grouped by PEtracer genotype calls. Genotype calls are first stratified by the PEtracer call (top two rows, last three rows), and further stratified by the LAML-Pro genotype call within each category. Each row reports the number of genotypes in this cross-classification and the distribution of PEtracer call probabilities in this category. (D) PEtracer-colonies. Siblings ‘cell-21930’ (cellA) and ‘cell-22314’ (cellB), paired in the LAML-Pro tree. First two rows show PEtracer genotype call probability (y-axis) over all sites where PEtracer calls different genotypes for the two cells (x-axis). Bars show the argmax genotype probability, with red dashed lines at 0.70 denote PEtracer call threshold. Solid bars are above the call probability threshold and unfilled bars are below the threshold. PEtracer calls the same genotype state in both cells, with the exception of site 24 (indicated by asterisk). LAML-Pro estimates same genotypes for both cells.

#### LAML-Pro resolves missing and uncertain imaging-based observations and genotypes

Next, we compare LAML-Pro and the PEtracer genotypes and trees on PEtracer-colonies, which are *in situ* 4T1 cells cultured on a coverslip over the course of 6 days and traced using the PEtracer system with imaging-based readout. Following the PEtracer study, we focus our analysis on Colony 3, which has 695 cells and 48 sites.

First comparing cell genotypes, we find that the PEtracer pipeline lacks genotype calls at a substantial fraction of the sites: 20.4% of sites are missing imaged observations. The PEtracer pipeline to call genotypes additionally removes classifications made with low posterior probability (≤ 70%), so that the fraction of missing observations increases to 22.3%. In contrast, LAML-Pro *completely eliminates missing genotypes* (Fig. 3B, missing data in gray). LAML-Pro infers dropout events at 20.6% of sites and accordingly infers genotypes. LAML-Pro imputes the remaining 1.7% of sites as silenced, inferring no observation. We observe no bias in inferred versus observed genotype frequencies (Supp. Fig. S12).

To evaluate the accuracy of the LAML-Pro inferred genotypes, we evaluated LAML-Pro and PEtracer on sites where both methods called genotypes. LAML-Pro and PEtracer genotypes agree on 97.9% of observed sites. We quantified the probability of the PEtracer genotype calls under the PEtracer discriminative classifier. When LAML-Pro calls genotypes differently from PEtracer (on 2.1% of all sites with observations), the corresponding PEtracer genotype calls were on average made with lower confidence relative to all PEtracer genotype calls (Fig. 3C, contrasting rows (1, 3) with rows (2, 4, 5).) On sites where both methods called genotypes as the “unedited” state, the PEtracer genotype was called with high confidence (mean call probability of 0.98). In contrast, on sites where PEtracer called “unedited” genotypes but LAML-Pro disagreed, the PEtracer genotype call had lower confidence (mean call probability of 0.88). Similarly, when LAML-Pro and PEtracer called the same “edited” genotype, the PEtracer genotype was called with relatively high confidence (mean call probability = 0.98). In contrast, PEtracer “edited” genotypes were called with much less confidence on sites where LAML-Pro called genotypes differently – either as “unedited” (mean call probability = 0.67) or as a different “edited” genotype (mean call probability = 0.71). On sites where PEtracer and LAML-Pro genotype calls agree, LAML-Pro calls genotypes with high confidence (mean LAML-Pro probability = 0.99 for sites called unedited by both PEtracer and LAML-Pro or sites having same edit by both methods). On sites where PEtracer and LAML-Pro genotype calls disagree, LAML-Pro genotype calls also show lower confidence (mean LAML-Pro probability; PEtracer unedited, LAML-Pro edited= 0.95; PEtracer edited, LAML-Pro different edit = 0.94; PEtracer edited, LAML-Pro unedited = 0.83).

A qualitative example where the LAML-Pro and PEtracer lineage and genotype calls differ occurs at the sibling pair (‘21930’, ‘22314’) in the LAML-Pro tree (phylogenetic distance 0.12); the pair (‘21930’, ‘22314’) are not siblings in the PEtracer tree (phylogenetic distance 2.24). Several sites in both cells have PEtracer classification probabilities which fall below the lineage marker threshold of 70%, and would therefore be excluded from the PEtracer genotype calls (Fig. 3D). Across the 27 sites where both leaves have observations, PEtracer calls 25 with identical state and LAML-Pro calls 24 with identical state. However, since PEtracer additionally discards genotype calls with probability lower than 70% before tree inference, the PEtracer tree reflects that this pair of leaves shares only 20 identical states. In contrast, the LAML-Pro tree has lower phylogenetic distance between the two leaves, reflecting that this pair shares consistently similar – albeit lower probability – observations for a total of 24 shared sites.

#### LAML-Pro reconstructs lineage trees with higher spatial concordance compared to PE-tracer trees

The imaging-based observations in PEtracer provide spatial coordinates (*x, y*) for each cell. The PEtracer publication [Koblan et al., 2025] leverages correlation between phylogenetic distance and spatial distance as an orthogonal metric to evaluate tree topology. Quantitatively, the LAML-Pro tree is substantially different from the published PEtracer tree (RF=0.54) and the LAML tree (RF=0.86). The tree inferred by LAML-Pro had higher correlation (Pearson’s *R* = 0.06, *p* = 6.8 × 10^−65^) compared to the published PEtracer tree (Pearson’s *R* = 0.0, *p* = 0.55) and equal correlation to the LAML tree (Pearson’s *R* = 0.06, *p* = 0.0) – see also Supp. Fig. S13. We also evaluated under a Brownian motion model of cellular migration (Supp. Sec. S2.4), and found that the LAML-Pro tree had higher likelihood than the PEtracer tree, although worse likelihood than the LAML tree with log-likelihood LAML-Pro = −6.2, PEtracer= −7.4, LAML = −5.8).

Next we validated the spatial concordance of the trees inferred by LAML-Pro on an additional 111 cells filtered out by the PEtracer pipeline due to genotype uncertainty. Specifically, the PEtracer pipeline filters out cells with *<* 60% cassette detection; we relax this threshold to 50%, gaining an additional 111 cells, and apply LAML-Pro to this superset of cells. The LAML-Pro tree on this larger set of cells maintains the spatial correlation (Pearson’s *R* = 0.06, *p* = 0.0) and further improves the log-likelihood of the ancestral migration history (LAML-Pro 806 cells (−5.88)).

#### LAML-Pro lineage tree improves concordance with static barcodes

To further evaluate LAML-Pro trees and illustrate the applicability to datasets readout by other modes, we next use the PEtracer-barcodes dataset with 3,108 4T1 cells readout by sequencing. LAML-Pro produced lineage trees which better grouped cells sharing static barcodes introduced at two timepoints, compared to the PEtracer tree. Quantitatively, LAML-Pro required fewer inferred integration events at both timepoints (see Supp. Sec S2.4.3).

### 3.3 LAML-Pro recovers spatial concordance on baseMEMOIR

We analyzed mouse embryonic stem cells (mESCs) lineage traced with the base editing lineage tracing system baseMEMOIR over the course of 72 hours [Chadly et al., 2024]. The cell line is engineered with 396 sites, with 4 possible states at each site (3 possible edit states and 1 unedited state), read out as 16 channel pixel intensities by multiplexed Zombie-FISH imaging [Chadly et al., 2024]. Focusing on the two largest colonies (2 and 5), we compare the genotypes and trees inferred by LAML-Pro to those reported in the baseMEMOIR manuscript [Chadly et al., 2024]. As baseMEMOIR reported ultrametric trees, we ran LAML-Pro (ultrametric).

#### LAML-Pro leverages informative observations dropped by baseMEMOIR

The baseMEMOIR pipeline drops genotype calls with low confidence, resulting in genotype calls at only half of target sites (51.6% in Colony 2 with 30 cells and 58.6% in Colony 5 with 28 cells). However, imaging observations exist at a large fraction of these dropped sites (Colony 2 = 38.2%, Colony 5 = 41.06%); LAML-Pro is able to leverage all such observations (total observed fraction; Colony 2 = 70.1%, Colony 5 = 75.6%) and call genotypes at all sites.

To evaluate the LAML-Pro genotypes, we first analyzed the sites where baseMEMOIR produced genotype calls (Colony 2=70.1%, Colony 5=75.6% of sites). The LAML-Pro and baseMEMOIR genotype calls agree on a majority of genotype calls (81.4% of observed sites in Colony 2, and 83.1% of observed sites in Colony 5). On genotype calls which agreed, the baseMEMOIR genotype was called with high confidence (mean call probability in Colony 2=0.89, and Colony 5=0.90; Fig. 4A; Supp. Fig. S17). In contrast, when LAML-Pro and baseMEMOIR disagreed on genotype calls (Colony 2=18.6%, Colony 5=16.9% of baseMEMOIR genotype calls), the baseMEMOIR genotype was called with low confidence relative to agreements (mean call probability; Colony 2=0.65, Colony 5=0.66). Most disagreements between baseMEMOIR and LAML-Pro are attributed to the case when baseMEMOIR called an “edited” state. In nearly all such disagreements, the baseMEMOIR genotype was called with confidence below the baseMEMOIR confidence threshold, so that these genotypes are not considered in the downstream baseMEMOIR tree inference (Fig. 4A). Finally, the LAML-Pro genotype frequencies on imputed genotypes are similar to the LAML-Pro genotype frequencies on sites with observations (Supp. Table S4), providing further credence to the LAML-Pro genotype calls.

**Figure 4.**
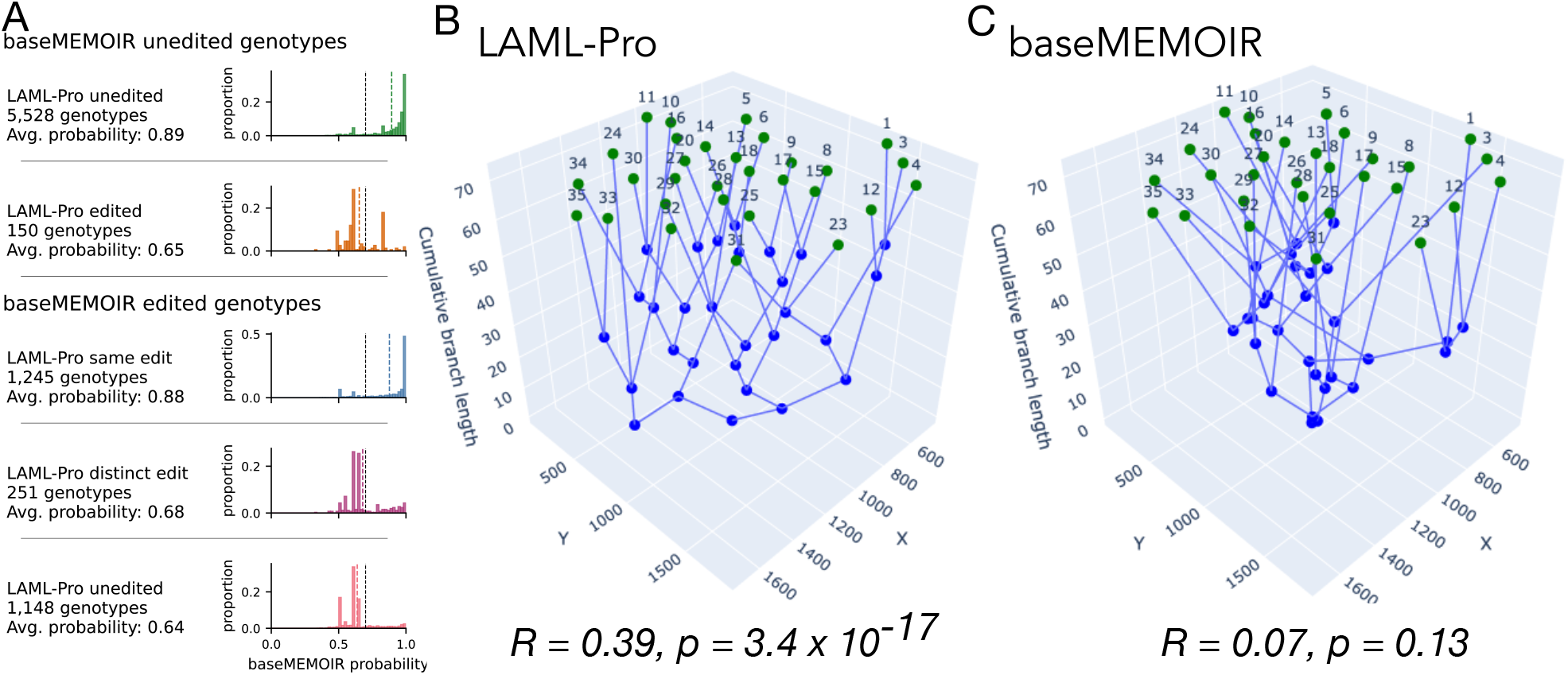
baseMEMOIR Colony 2. (A) Genotype differences between LAML-Pro and baseMEMOIR, grouped by baseMEMOIR genotype calls. Genotypes are first stratified by the baseMEMOIR call (top two rows, last three rows), and further stratified by the LAML-Pro genotype call within each category. Each row reports the number of genotypes in this cross-classification and the distribution of baseMEMOIR call probabilities in this category. (B) The inferred LAML-Pro tree and (C) baseMEMOIR tree visualized in 3D: (*x, y*) are given by imaged spatial location and *z* is given by the cumulative estimated branch length. *R* indicates Pearson correlation coefficient, and *p* indicates corresponding p-value.

#### LAML-Pro tree improves genotype concordance with baseMEMOIR genotypes

To characterize the substantial difference between the LAML-Pro and baseMEMOIR trees (Colony 2 RF=0.96, Colony 5 RF=1.0), we evaluate the trees in terms of agreement with the baseMEMOIR genotypes. Although the advantage is to baseMEMOIR trees in this evaluation, LAML-Pro trees achieve substantially higher genotype concordance by several metrics. Given a pair of cells, we compute the expected Hamming distance (EHD) between their corresponding baseMEMOIR genotype probabilities. We compare the EHD for all sibling pairs (evaluation metrics in Supp. Sec. S2.3.1), and find that LAML-Pro trees consist of siblings with much lower median normalized EHD than the baseMEMOIR trees (Colony 2: LAML-Pro=0.094, baseMEMOIR=0.348; Colony 5: LAML-Pro=0.121, baseMEMOIR=0.762; Supp. Figs. S18). For nearly every leaf, the sibling in the LAML-Pro topology had better genotype concordance than the corresponding baseMEMOIR sibling (Supp. Fig. S19).

#### LAML-Pro tree improves spatial concordance compared to the baseMEMOIR tree

We evaluate the LAML-Pro, LAML and baseMEMOIR trees in terms of two metrics of spatial concordance. Although the baseMEMOIR trees were inferred under a Bayesian phylogeography model which takes the spatial coordinates of each cell as additional input (Supp. Sec. S2.5.2), the phylogenetic distances from the LAML-Pro tree have greater correlation with the spatial Euclidean distances on both Colony 2 (LAML-Pro *R* = 0.349, *p* = 3.4 × 10^−17^; LAML *R* = 0.003, *p* = 0.95; baseMEMOIR *R* = −0.073, *p* = 0.13) and Colony 5 (LAML-Pro *R* = 0.229, *p* = 1.5 × 10^−5^; LAML *R* = 0.195, *p* = 2.4 × 10^−4^; baseMEMOIR *R* = 0.003, *p* = 0.96) (Fig. 4B, C; Supp. Fig. S17). The phylogeography model under which the baseMEMOIR trees are inferred makes an implicit assumption of cell migration under Brownian motion [Chadly et al., 2024, Bouckaert, 2016]. Under the same assumption of Brownian motion migration, we compute the maximum likelihood ancestral migration history on each topology. We observe that the LAML-Pro tree supports a higher likelihood ancestral migration than the baseMEMOIR tree in both Colony 2 (log-likelihood LAML-Pro = −6.5, LAML= −6.6, baseMEMOIR= −6.8) and in Colony 5 (log-likelihood LAML-Pro = −6.7, LAML= −6.6, baseMEMOIR= −6.9). We use the inferred spatial migration (Supp. Sec. S2.5.4) to reconstruct the LAML-Pro and baseMEMOIR trees in 3D (Fig. 4B, C). We note that the baseMEMOIR trees have long branch lengths at the leaves (Fig. 4C). The LAML-Pro tree has lower branch length variance across the tree than the baseMEMOIR trees, on Colony 2 (LAML-Pro =225.4, baseMEMOIR=273.3) and Colony 5 (LAML-Pro =46.33, baseMEMOIR=127.7).

### 3.4 LAML-Pro infers trees quickly and scalably

Despite the more complex observation model, LAML-Pro runtime remains comparable to a re-implementation of LAML in C++. For example, the median time taken by LAML-Pro to infer a cell lineage tree with *n* = 300 simulated cells using a fixed number of 5,000 proposed NNI moves is 1 hour and 42 minutes compared to 1 hour and 39 minutes for our reimplementation of LAML. The runtime scales linearly with the number of cells for a fixed number of NNI iterations (Supp. Fig. S6). All methods were given a single core CPU.

Directly taking in the observations and handling genotype uncertainty does not increase LAML-Pro runtime. On 695 cells traced by PEtracer, our new implementation of LAML produced trees in 46 minutes on average, whereas LAML-Pro produced trees in 42 minutes on average (each given 5,000 moves). For comparison, LAML-Pro was given 25,000 moves on a second PEtracer experiment with 3,108 cells, which took 18 hours. On baseMEMOIR LAML-Pro converged on trees quickly (Colony 2 with 30 cells in 6 minutes and Colony 5 with 28 cells in 4.6 minutes), whereas LAML took slightly longer (Colony 2 and 5 both took 13 minutes on average, each given 5,000 moves). LAML-Pro and LAML were run with the ultrametric constraint to match the original papers’ trees [Koblan et al., 2025, Chadly et al., 2024], and given twelve initial trees. LAML-Pro improved all initial trees into nearly the same topology (Supp. Fig. S20B).

## 4 Discussion

In this paper, we introduced LAML-Pro, a method to simultaneously infer the cell lineage tree and cell genotypes directly from imaging or sequencing-based DLT observations without first calling genotypes. There are several future directions. First, the procedures to process DLT data vary across studies, and the heuristics used in these processing steps might be reevaluated in light of LAML-Pro’s ability to incorporate data that would have previously been discarded as low confidence. Second, assessing the statistical confidence of branches (e.g. using the bootstrap) would be helpful for interpretation and downstream analyses. Third, the PMMO model framework easily extends to accommodate challenges specific to new lineage tracing technologies, such as sequencing-based observations convolving 1-3 cells [Jones et al., 2024]. Similarly, the incorporation of priors in genotype inference to model known sources of genotype conflict, such as persisting transcripts [Koblan et al., 2025], or to leverage data with heterogeneous sampling times, may further improve reconstruction and branch length accuracy. Fourth, evaluating the validity of the ultrametric constraint on particular biological datasets would be helpful. Finally, LAML-Pro consistently scales to thousands of cells in existing experimental datasets. Additional scaling for future whole organ/organism lineage tracing datasets may require additional techniques such as including SPR and TBR tree search moves [Swofford, 1990], parallelization [Ott et al., 2007], divide-and-conquer techniques [Warnow, 2019, Molloy et al., 2019], or constraint clades [Dai et al., 2025]).

## 6 Acknowledgments

We thank Uyen Mai for helpful discussions and the anonymous reviewers for valuable feedback.

