## Supplementary Materials for "LAML-Pro: Joint Maximum Likelihood Inference of Cell Genotypes and Cell Lineage Trees"

### S1. Supplementary Methods

#### S1.1. PMMO model

On a single edge $e$ at target site $k$, the transition probability matrix $\mathbf{P}$ can be computed as $\mathbf{P}_\delta^{\mathbf{Q}} = \mathrm{e}^{\mathbf{Q}\delta}$. Let $\delta = \lambda t$ be the branch length of edge $e$ in mutation units. For the PMMO model, the matrix exponential has the following closed form,

$$\mathrm{e}^{\mathbf{Q}\delta} = \begin{array}{c} \\ 0 \\ 1 \\ \vdots \\ M \\ -1 \end{array} \begin{array}{c} \begin{array}{cccccc} 0 & 1 & \dots & M & -1 \end{array} \\ \begin{bmatrix} e^{-\delta(1+\nu)} & q_1 e^{-\delta\nu}(1-e^{-\delta}) & \dots & q_m e^{-\delta\nu}(1-e^{-\delta}) & 1-e^{-\delta\nu} \\ 0 & e^{-\delta\nu} & \dots & 0 & 1-e^{-\delta\nu} \\ \vdots & \vdots & \ddots & \vdots & \vdots \\ 0 & 0 & \dots & e^{-\delta\nu} & 1-e^{-\delta\nu} \\ 0 & 0 & \dots & 0 & 1 \end{bmatrix} \end{array}. \tag{S1}$$

#### S1.2. Generating sequencing-based observations.

To determine the observation model density $\Gamma(x|z)$ in Eq. 3, it suffices to specify the conditional density $f_z(x)$ of generating a set of reads $x$ from the hidden state $z$. We model $f_z(x)$ as a multinomial distribution, with $n_u$ independent draws and probabilities $\{p_s^z\}$ for drawing a state $s$ read given hidden state $z$.

For each cell $u$ we observe $n_u$ reads each taking one of $d$ states. Our observation alphabet is $\{x(u) \in \mathbb{N}_0^d : \sum_{s=1}^{d} x_s(u) = n_u\} \cup \{?\}$, where each observation $x_s(u)$ denotes the number of state $s$ reads in cell $u$ and summing over the states yields cell $u$'s total number of reads $n_u$. Let $m(s) = \sum_u x_s(u)$ be the total expression of state $s$. Let $\rho$ be the probability of correct assignment between UMI and cell. To match biology, we model higher probability of incorrectly sampling transcripts which are overexpressed, so that:

$$p_s^z := \begin{cases} \rho, & \text{if } s = z, \\ (1-\rho)\frac{m(s)}{\sum\limits_{s' \neq z} m(s')} & \text{otherwise,} \end{cases} \tag{S2}$$

Many sources of error may be present in sequencing-based observations (e.g. PCR errors, sequencing errors, ambient RNA) [Jones et al., 2020, Koblan et al., 2025]. Reads are filtered with a series of reasonable heuristics, before the (hidden) character state for each cell and site is determined by taking whichever genomic edit has the most supporting transcripts [Chan et al., 2019, Jones et al., 2020]. Above we defined $f_z$ to model the generation of common ambient RNA errors, which come from mis-attributing over-expressed states with many reads to cells during the single cell sequencing process (Supplementary Figure S7).

#### S1.3. Generating a character matrix observation.

We take the PMM model [Chu et al., 2025] as our representative model assuming the observed data is the character matrix. Under the PMM model, the observed alphabet consists of states $\{0, 1, \dots, M, ?\}$. Since the observation alphabet is a discrete set, in this model the observed data is compactly represented by an $N \times K$ character matrix, where $N$ is the number of observed cells and $K$ is the number of sites. The observation model is implicit; given an $N \times K$ PMM character matrix [Chu et al., 2025], the observation model is governed only by the dropout probability $\phi$. That is, defining $f_z(x) = \mathbf{1}[z = x]$ recovers the original PMM model, which does not allow for any errors in the character matrix. This assumption oversimplifies the data, as the presence of errors in the character matrix is documented and nontrivial to correct Jones et al. [2020].

A simple extension describes errors in the character matrix. Let $0 \leq \eta \leq 1 - \frac{1}{M}$ denote the error probability. We specify the observation model by defining

$$f_z(x) := \begin{cases} \frac{1-\phi-\eta}{1-\phi} & \text{if } z = x, \\ \frac{\eta}{M(1-\phi)} & \text{otherwise.} \end{cases} \tag{S3}$$

This formulation is analogous to error channels in coding theory. Without missing data, our formulation corresponds to a discrete memoryless $M$-ary symmetric channel in which each transmitted symbol $z$ (from alphabet of size $M$) might be incorrectly received with probability $\eta$ (i.e. the "crossover probability" [Shannon, 1948, Gallager, 1968]). The model of missing data corresponds to an erasure channel, where each symbol is dropped with probability $\phi$ [Gallager, 1968].

#### S1.4. Maximum likelihood estimation under the PMMO model

To perform maximum likelihood estimation of the parameters $\Theta$ on a fixed tree $\mathcal{T}$, we use an expectation maximization algorithm defined via the update in Eq. 5. Let $\Omega = \{0, 1, \ldots, M, -1\}$ denote the hidden alphabet, $\pi$ be the probability distribution of the root $r_T$, and $\Theta^t$ denote the estimated parameters at iteration $t$ of the algorithm. The form of the expected complete log-likelihood $Q(\Theta \mid \Theta^t)$ in Eq. 6 is derived following the argument in Siepel [2002] and Chu et al. [2025].

To derive the form of $Q(\Theta \mid \Theta^t)$ in Eq. 6, we first expand $Q(\Theta \mid \Theta^t)$ and group like terms, obtaining

$$\begin{aligned} Q(\Theta \mid \Theta^t) &= \mathbb{E}_{Z \sim \mathbb{P}(\cdot \mid X, \mathcal{T}, \Theta^t)}[\log \mathbb{P}(X, Z \mid \mathcal{T}, \Theta)] \\ &= \sum_z \mathbb{P}(Z = z \mid X, \mathcal{T}, \Theta^t) \cdot \log \mathbb{P}(X = x, Z = z \mid \mathcal{T}, \Theta) \\ &= \sum_z \mathbb{P}(Z = z \mid X, \mathcal{T}, \Theta^t) \cdot \log \Big[\pi_{z(r)} \\ &\qquad \cdot \prod_{e=(u,v)} \mathbf{P}^{\mathbf{Q}}_{z(u),z(v),\delta_e} \cdot \prod_{u \in V_L} \mathbf{\Gamma}(x(u) \mid z(u))\Big] \\ &= \sum_z \mathbb{P}(Z = z \mid X, \mathcal{T}, \Theta^t) \cdot \Big[\log \pi_{z(r)} \\ &\qquad + \sum_{e=(u,v)} \log \mathbf{P}^{\mathbf{Q}}_{z(u),z(v),\delta_e} + \sum_{u \in V_L} \mathbf{\Gamma}(x(u) \mid z(u))\Big]. \end{aligned} \tag{S4}$$

Next, we analyze the three logarithmic terms in Eq. S4 using the strategy described in Siepel [2002]. First, the term corresponding to the root state $z_r$ is equal to

$$\begin{aligned} &\sum_z \mathbb{P}(Z = z \mid X, \mathcal{T}, \Theta^t) \cdot \log \pi_{z(r)} \\ &= \sum_{a \in \Omega} \mathbb{P}(Z(r_T) = a \mid X, \mathcal{T}, \Theta^t) \cdot \log \pi_a. \end{aligned} \tag{S5}$$

This follows by marginalizing the hidden states at all nodes $u \neq r_T$. However, since the root distribution $\pi$ is assumed to be fixed, this is a constant term and is independent of the solution to the optimization problem in the Mstep.

Reordering the sums, introducing an indicator variable for the transition along each edge $e$, applying a marginalization argument, and using the definition of $\mathcal{S}_{\mathsf{edge}}$, we obtain the equality

$$\begin{aligned} &\sum_z \sum_{e=(u,v)} \mathbb{P}(Z = z \mid X, \mathcal{T}, \Theta^t) \cdot \log \mathbf{P}^{\mathbf{Q}}_{z(u),z(v),\delta_e} \\ &= \sum_{e=(u,v)} \sum_z \sum_{a,b \in \Omega} \mathbb{P}(Z = z \mid X, \mathcal{T}, \Theta^t) \cdot \log \mathbf{P}^{\mathbf{Q}}_{a,b,\delta_e} \\ &\qquad \cdot \mathbf{1}(z(u) = a, z(v) = b) \\ &= \sum_{e=(u,v)} \sum_{a,b \in \Omega} \log \mathbf{P}^{\mathbf{Q}}_{a,b,\delta_e} \cdot \sum_z \mathbb{P}(Z = z \mid X, \mathcal{T}, \Theta^t) \\ &\qquad \cdot \mathbf{1}(z(u) = a, z(v) = b) \\ &= \sum_{e=(u,v)} \sum_{a,b \in \Omega} \mathcal{S}_{\mathsf{edge}}(e, a, b \mid \mathcal{T}, \Theta^t) \cdot \log \mathbf{P}^{\mathbf{Q}}_{a,b,\delta_e} \end{aligned}$$

for the second logarithmic term in Eq. S4.

Following an identical argument, we obtain the equality

$$\sum_{z}\sum_{u\in V_L}\mathbb{P}(Z=z\mid X,\mathcal{T},\Theta^t)\cdot\log\mathbf{\Gamma}(x(u)\mid z(u)) = \sum_{u\in V_L}\sum_{a\in\Omega}\mathcal{S}_{\mathsf{node}}(u,a\mid\mathcal{T},\Theta^t)\cdot\log\mathbf{\Gamma}(x(u)\mid a) \tag{S6}$$

for the third logarithmic term in Eq. S4.

Adding the terms Eq. S5-S6 together results in the formula

$$\begin{aligned} Q(\Theta\mid\Theta^t) = &\sum_{e=(u,v)}\sum_{a,b\in\Omega}\mathcal{S}_{\mathsf{edge}}(e,a,b\mid\mathcal{T},\Theta^t)\cdot\log\mathbf{P}^{\mathbf{Q}}_{a,b,\delta_e} \\ &+\sum_{u\in V_L}\sum_{a\in\Omega}\mathcal{S}_{\mathsf{node}}(u,a\mid\mathcal{T},\Theta^t)\cdot\log\mathbf{\Gamma}(x(u)\mid a) \\ &+\sum_{a\in\Omega}\mathbb{P}(Z(r_T)=a\mid X,\mathcal{T},\Theta^t)\cdot\log\pi_a \end{aligned} \tag{S7}$$

which, ignoring the constant term from Eq. S5, completes the derivation of $Q(\Theta\mid\Theta^t)$ stated in Section 2.2.

##### S1.4.1. M-Step

The M-step maximizes the function $Q(\Theta\mid\Theta^t)$ with respect to the $\Theta$ subject to the constraints in Eq. 1. Let $\mathcal{K}$ denote the convex set of constraints in Eq. 1. In this section, we describe the details of this optimization problem and the steps taken to efficiently solve it.

First, we expand the observation term in Eq. S6, obtaining

$$\begin{aligned} &\sum_{u\in V_L}\sum_{a\in\Omega}\mathcal{S}_{\mathsf{node}}(u,a\mid\mathcal{T},\Theta^t)\cdot\log\mathbf{\Gamma}(x(u)\mid a) \\ &=\sum_{u\in V_L}\sum_{a\neq -1}\mathcal{S}_{\mathsf{node}}(u,a\mid\mathcal{T},\Theta^t)\cdot[\log\phi\cdot\mathbf{1}(x(u)=?) \\ &\quad+(\log(1-\phi)+\log f_a(x(u)))\cdot\mathbf{1}(x(u)\neq ?)] \\ &\quad+\sum_{u\in V_L}\mathcal{S}_{\mathsf{node}}(u,a\mid\mathcal{T},\Theta^t)\cdot\mathbf{\Gamma}(x(u)\mid -1). \end{aligned} \tag{S8}$$

The terms $\mathcal{S}_{\mathsf{node}}(u,a\mid\mathcal{T},\Theta^t)\cdot\mathbf{\Gamma}(x(u)\mid -1)$ and $\mathcal{S}_{\mathsf{node}}(u,a\mid\mathcal{T},\Theta^t)\cdot\log f_a(x(u)))$ in the preceding are constant with respect to the optimization variable $\Theta$ and can be ignored during the M-step.

Second, we note that, due to the structure of the PMMO model Chu et al. [2025], $\mathbf{P}^{\mathbf{Q}}_{a,b,\delta}$ have the closed forms obtained by exponentiating the transition rate matrix in Eq. (2):

$$\log\mathbf{P}^{\mathbf{Q}}_{0,0,\delta} = -\delta(1+\nu), \tag{S9}$$

$$\log\mathbf{P}^{\mathbf{Q}}_{-1,-1,\delta} = 0, \tag{S10}$$

$$\log\mathbf{P}^{\mathbf{Q}}_{0,a,\delta} = \log q_a - \delta\nu + \log(1-e^{-\delta}) \quad \text{for } a\neq -1, \tag{S11}$$

$$\log\mathbf{P}^{\mathbf{Q}}_{a,-1,\delta} = \log(1-e^{-\delta\nu}) \quad \text{for } a\neq -1, \tag{S12}$$

$$\log\mathbf{P}^{\mathbf{Q}}_{a,a,\delta} = -\delta\nu \quad \text{for } a\neq 0,-1. \tag{S13}$$

For the remaining $a, b \in \Omega$, $\log \mathbf{P}^{\mathbf{Q}}_{a,b,\delta} = -\infty$. Then, we define the six quantities

$$\begin{aligned}
\mathcal{C}_e^{z\to z} &= \mathcal{S}_{\mathsf{edge}}(e, 0, 0 \mid \mathcal{T}, \Theta^t), \\
\mathcal{C}_e^{z\to a} &= \sum_{b\neq -1} \mathcal{S}_{\mathsf{edge}}(e, 0, b \mid \mathcal{T}, \Theta^t), \\
\mathcal{C}_e^{a\to m} &= \sum_{b\neq -1} \mathcal{S}_{\mathsf{edge}}(e, b, -1 \mid \mathcal{T}, \Theta^t), \\
\mathcal{C}_e^{a\to a} &= \sum_{b\neq 0,-1} \mathcal{S}_{\mathsf{edge}}(e, b, b \mid \mathcal{T}, \Theta^t), \\
\mathcal{C}_m &= \sum_{u\in V_L} \sum_{a\neq -1} \mathbf{1}(x(u) = ?) \cdot \mathcal{S}_{\mathsf{node}}(u, a \mid \mathcal{T}, \Theta^t), \text{ and} \\
\mathcal{C}_n &= \sum_{u\in V_L} \sum_{a\neq -1} \mathbf{1}(x(u) \neq ?) \cdot \mathcal{S}_{\mathsf{node}}(u, a \mid \mathcal{T}, \Theta^t),
\end{aligned} \tag{S14}$$

which are constant for fixed $\Theta^t$. It follows by ignoring constant terms in the optimization problem that

$$\begin{aligned}
&\arg\max_{\Theta\in\mathcal{K}} Q(\Theta \mid \Theta^t) \\
&= \arg\max_{\Theta\in\mathcal{K}} \Bigg\{ \sum_{e=(u,v)} \big[ -\delta_e(1+\nu)\cdot \mathcal{C}_e^{z\to z} \\
&\qquad + (\log(1-e^{-\delta_e}) - \delta_e\nu)\cdot \mathcal{C}_e^{z\to a} \\
&\qquad + \log(1-e^{\delta_e\nu})\cdot \mathcal{C}_e^{a\to m} - \delta_e\nu\cdot \mathcal{C}_e^{a\to a}\big] \\
&\quad + \log(\phi)\cdot \mathcal{C}_m + \log(1-\phi)\cdot \mathcal{C}_n \Bigg\}.
\end{aligned} \tag{S15}$$

As a function of the inferred parameters $\delta_e, \nu$, and $\phi$, Eq. S15 is a non-convex optimization problem subject to convex constraints. However, there are several properties of Eq. S15 that make it amenable to efficient numerical optimization. First, since there is no dependence between $\delta_e$ and $\delta_{e'}$ with $e \neq e'$ in the objective of Eq. S15, all second-order partial derivatives $\frac{\partial}{\partial\delta_e\partial\delta_{e'}}$ are zero. Consequently, the Hessian of the objective in Eq. S15 contains $\mathcal{O}(N+M)$ non-zero entries where $N$ is the number of branches and $M$ is the alphabet size. This enables the efficient application of second-order methods to solve the optimization problem in Eq. S15 as Hessian-vector products can be computed in $\mathcal{O}(N+M)$ time. Second, the parameter $\phi$ is separable and can be optimized independently of $\nu$ and $\delta_e$. Finally, the gradient of the objective of Eq. S15 has a simple and easily computable closed form.

To perform the optimization, we use an interior point method; this is a standard class of second-order algorithms for constrained, non-convex optimization Nocedal et al. [2006]. Specifically, we use the algorithm detailed in Wächter et al. [2006] and implemented in the open-source package IPOPT Biegler et al. [2009]. The key advantage of using an interior point method compared to a first-order algorithm, such as projected gradient descent Nocedal et al. [2006], is twofold. The interior point method obtains i) fast (i.e. quadratic) local convergence and ii) is guaranteed to converge to a stationary (KKT) point. Further, since the Hessian is sparse, the increase in iteration complexity of using a second-order method as opposed to a first-order method is slight.

###### S1.4.2. Topology search

### S2. Supplementary Results

#### S2.1. Evaluation metrics

**The normalized Robinson-Foulds (RF) distance.** Given two trees $T_1, T_2$, we measure the topological distance between the two trees with the Robinson-Foulds (RF) distance, defined as

$$\Delta_{RF}(T_1, T_2) = \frac{FN + FP}{n_{\text{internal}}(T_1) + n_{\text{internal}}(T_2)}. \tag{S16}$$

**Algorithm 1** Pseudocode for LAML-Pro. Inputs: observations $x$, starting tree $\mathcal{T}_0$. Hyperparameters: maximum iterations $t_{\max}$ (default = 20000), simulated annealing parameters defaulting to $T_{\text{init}} = 0.01, \alpha = 0.99$.

**function** LAML-PRO($x$, $\mathcal{T}_0$)
  $\mathcal{T} \leftarrow \mathcal{T}_0$
  $\hat{\Theta}' \leftarrow \arg\max_{\Theta} \log L(\mathcal{T}, \hat{\Theta}; x)$ ▷ EM algorithm
  $c \leftarrow 0$ ▷ number of accepts
  **for** $t = 1, \ldots, t_{\max}$ **do**
    $\mathcal{T}' \leftarrow$ sample new topology via random NNI of $\mathcal{T}$
    $\hat{\Theta}' \leftarrow \arg\max_{\Theta} \log L(\mathcal{T}', \hat{\Theta}; x)$ ▷ EM algorithm
    $\Delta \leftarrow \log L(\hat{\Theta}'; \mathcal{T}'; x) - \log L(\hat{\Theta}; \mathcal{T}; x)$
    $p \leftarrow e^{\Delta/(T_{\text{init}} \cdot \alpha^c)}$
    **if** $\Delta > 0$ or $D \sim \text{Unif}(0,1) \leq p$ **then** ▷ simulated annealing
      $\mathcal{T} \leftarrow T'$.
      $c \leftarrow c + 1$.
    **end if**
  **end for**
  **return** $\mathcal{T}, \hat{\Theta}$
**end function**

To compute the RF distance on the same leafset, we compute the false positives (bipartitions in $T_2$ not present in $T_1$ and false negatives (bipartitions in $T_1$ not present in $T_2$). We normalize the RF distance by the total number of internal edges in both threes given by $n_{\text{internal}}$.

**Phylogenetic distance.** The phylogenetic distance between two nodes in a tree is defined as the sum of branch lengths along the unique path connecting them in the tree.

**Spatial Euclidean distance.** Given two pairs of points with spatial coordinates in 2D: $(a, b)$ and $(c, d)$, the spatial Euclidean distance is given by $d = \sqrt{(-a)^2 + (d - b)^2}$.

#### S2.2. Simulated data

A model tree topology $T$ with 4,096 leaves is simulated using a pure birth model Jones et al. [2020], with branch lengths $\{\delta_e\}_{e \in T}$ are sampled independently from a lognormal distribution ($\mu = 1, \sigma = 0.1$). Hidden sequences ($K = 400$) evolve i.i.d. down the model tree under the PMMO model with a hidden alphabet of four states $\{0, 1, 2, 3\}$ and a uniform transition rate matrix. To obtain $\approx 40\%$ edited (non-zero) entries in simulated data we set the editing rate $\lambda = 0.095$ and make the molecular clock assumption.

We construct a mixture of 4 isotropic multivariate Gaussians in 3 dimensions, where the observation corresponding to hidden state $s \in \mathcal{A}$ is sampled from $\mathcal{N}(\mu_s, \sigma^2 I)$ (Fig. 2A). Gaussian means $\mu_s$ are fixed to the vertices of the regular tetrahedron with unit edge lengths (Fig. 2A). More explicitly, each $\mu_s$ is given by,

$$\mu_0 = \begin{bmatrix} -0.612 \\ 3.66 \times 10^{-17} \\ 0 \end{bmatrix}, \mu_1 = \begin{bmatrix} 0.204 \\ -0.577 \\ -1.38 \times 10^{-17} \end{bmatrix},$$
$$\mu_2 = \begin{bmatrix} 0.204 \\ 0.289 \\ -0.500 \end{bmatrix}, \mu_3 = \begin{bmatrix} 0.204 \\ 0.289 \\ 0.5 \end{bmatrix}.$$

We vary the shared standard deviation $\sigma$ in order to control the separation between each pair of distributions (Table S1). We additionally quantify the pairwise distribution separation using the Bhattacharya coefficient $\rho$ Bhattacharyya [1943],

$$\rho = \exp\left(-\frac{\|\boldsymbol{\mu}_1 - \boldsymbol{\mu}_2\|^2}{8\sigma^2}\right)$$

In total, we have 8 model conditions with varying missing and observation error. Including the sampling conditions, there are 24 total model conditions (Table S1), each with 5 sampling replicates, for a total of 120 replicates.

##### S2.2.1. Benchmarking against other methods

Using this simulated dataset, we benchmarked two ways of running LAML-Pro against two ways of running Neighbor Joining, as well as four ways of running LAML: LAML-Pro (ultrametric), LAML-Pro (No Ultra), NJ (HD), NJ (wHD), LAML (ultrametric), LAML (No Ultra), LAML (Ultra, True CM), LAML (No Ultra, True CM). To run LAML-Pro, we generate 11 starting trees: one neighbor joining tree and ten parsimony trees via random stepwise addition, using the 'phangorn' package Schliep [2011]. We provide this script to generate the starting trees in the '/scripts' folder on the LAML-Pro github repository.

Neighbor Joining was run using the implementation in 'biopython' [Cock et al., 2009]. We compute pairwise distances between cell genotypes using (i) the standard Hamming distance definition, and (ii) the weighted Hamming distance described in [Jones et al., 2020], which treats missing states as uninformative and skips all such sites. In our setting with heritable missing, this latter weighted metric performed surprisingly poorly. We found that normal Hamming Distance produced downstream trees with vastly improved tree reconstruction error. This is especially true when there are high rates of heritable missing, and is notable because the weighted Hamming distance is often used to generate the Neighbor Joining tree [Koblan et al., 2025]. Thus, we report results from normal Hamming Distance (HD) and weighted Hamming distance (wHD) in both the main text and Supplementary Figures.

In our evaluation of branch length accuracy, we additionally benchmarked our two ways of running LAML-Pro against ConvexML [Prillo et al., 2025]. We installed ConvexML version 1.0.0 from pip, and ran it as follows:

```
res = convexml.convexml(tree_newick_str, leaf_sequences)
```

Please note that ConvexML failed to produce results on 10 instances of the NJ (wHD) tree, in the missing data model condition. We turned multifurcations off in order to obtain results on the full set of simulations.

##### S2.2.2. Discussion of the simulation results

We observe that on average all methods achieve lower topology reconstruction error when there is no missing data than in the presence of high rates of missing data (Supp. Fig. S2). This trend is consistent across larger numbers of cells, and we observe that error increases as observations from one state are more easily mistaken for another state (i.e. as standard deviation $\sigma$ increases, see Supp. Fig. S2). Notably, we observe that NJ (wHD) performs worse than NJ (HD) even at low rates of confusion ($\sigma = 0.3, 0.4$) regardless of the presence of missing data. We observe that NJ (wHD) performs substantially worse than NJ (HD) at high confusion rates ($\sigma = 0.5$) in the presence of missing data. We observe that LAML-Pro (ultrametric) achieves lower topology reconstruction accuracy than LAML-Pro (No Ultra); this is perhaps not surprising as the data is simulated under an ultrametric model. However, this analysis does demonstrate that LAML-Pro is robust to such model misspecification (in this direction; i.e. specifying no ultrametric constraint), LAML-Pro is able to identify a close topology. Additional work is necessary to demonstrate the inverse.

Across $n = 100, 200, 300$ cells and confusion, we observe that LAML-Pro has substantially lower genotype error than the baseline approach of taking the genotype state with the argmax probability (given by Argmax CM, see Supp. Fig. S3). The margin of improvement increases as the observation confusion (given by the observation standard deviation $\sigma$) increases.

We find that LAML-Pro (ultrametric) and LAML-Pro (No Ultra) produce very similar parameter estimates for heritable silencing rate *nu* and dropout probability *phi* (Supp. Fig. S4. We observe that both ways of running LAML estimate both $\nu$ and $\phi$ with increasing error as the observations become more noisy ($\sigma$ increases). Notably, we observe that LAML-Pro (No Ultra) produces slightly more accurate estimates for $\nu$ at $n = 100$ and $n = 300$ cells. The ultrametric constraint does not affect the estimates of $\phi$ across the model conditions.

Finally, we observe that across the number of cells $N$ and the missing data conditions, LAML-Pro (ultrametric) infers *extremely accurate branch lengths* (Supp. Fig. S5). Despite some model misspecification, LAML-Pro (No Ultra) still estimates very accurate branch lengths. Both are comparable with LAML (True

CM), which is a difficult baseline to beat. As expected, the branch length error increases as the observation confusion increases ($\sigma$ increases).

#### S2.3. Evaluation Metrics on two lineage tracing systems

##### S2.3.1. Evaluation Metrics for Genotypes

To evaluate a tree topology's concordance with the cells' genotypes, we define several metrics below. For each leaf $c$, the tree topology defines a closest leaf $\text{sib}_T(c)$. We compute the expected hamming distance (or one of the variants below) between the genotypes of $c$ and $\text{sib}_T(c)$.

We normalize the three expected Hamming distance metrics below using the minimum pairwise distance between all pairs of genotypes $d_{\min}$ and maximum pairwise distances between all pairs of genotypes $d_{\max}$. We perform this normalization once, using the median $d_{\min}, d_{\max}$ to normalize the median $d$ for each metric:

$$\frac{d - d_{\min}}{d_{\max} - d_{\min}}$$

**Expected Hamming Distance between Genotypes (ignoring missing)** Below we define the expected Hamming distance (also referenced elsewhere as the "expected genotype disagreement" Li et al. [2009]).

Let $g(c)$ be the genotype vector of a leaf $c$, so that $g(c) = (G_1, G_2, G_3, \ldots, G_n)$. Each element $G_i$ is a genotype distribution, either (1) the observed categorical distribution over states $k \in \Sigma$, or (2) missing "?". Then over $n$ sites, let $\delta_i = 1$ if $P_i, Q_i$ are distributions, otherwise $\delta_i = 0$. Then we have that the "ignore-missing" expected hamming distance between two cells (ignoring all sites with missing data in either cell) is:

$$imEHD(P, Q) = \frac{1}{\sum\limits_i \delta_i} \sum_{i=1}^{n} \delta_i \left(1 - \sum_{k \in \Sigma} P_i(k) Q_i(k)\right)$$

**Expected Hamming Distance between Genotypes (missing-impute)** Since we have a significant number of sites with missing data, we define a second version of expected Hamming distance. Instead, let each element $G_i$ be a genotype distribution, either (1) the observed categorical distribution over states $k \in \Sigma$, or (2) in the case of a missing observation "?", use the empirical genotype distribution over all cells with observations at site $i$. Thus, over $n$ sites, we define the expected Hamming distance (EHD) as

$$EHD(P, Q) = \frac{1}{n} \sum_{i=1}^{n} \left(1 - \sum_{k \in \Sigma} P_i(k) Q_i(k)\right)$$

**Expected Hamming Distance between Genotypes (shared-missing)** Since the presence of missing data can also indicate the presence of a shared ancestral silenced state, the EHD may be over-penalizing sites where both cells' genotypes are missing data due to the high uncertainty in the empirical genotype distribution. Thus, the shared-missing expected Hamming distance captures the intuition that a pair of sites with no observations is still agreement between observations.

Let $g(c)$ be the genotype vector of a leaf $c$, so that we have $g(c) = (G_1, G_2, \ldots, G_n)$ as a vector of genotype distributions. Each element $G_i$ is a genotype distribution which is either observed or missing (filled in with the empirical genotype distribution over all cells with observations at site $i$) as above. We begin by defining the agreement between two genotype distributions,

$$A_i(P_i, Q_i) = \begin{cases} \sum\limits_{k \in \Sigma} P_i(k) Q_i(k) \text{ if both are observed,} \\ \sum\limits_{k \in \Sigma} \hat{p}_i(k) Q_i(k) \text{ if WLOG } P_i \text{ is missing, } Q_i \text{ is observed,} \\ \frac{1}{2} + \frac{1}{2} \sum_{k \in \Sigma} \hat{p}_i(k)^2 \text{ if both are missing} \end{cases}$$

Note that in this last case, the two cases are agreement due to the shared silenced state, and agreement due to shared dropout. This assumes that both types of missing are equally likely. Over $n$ sites, the shared-missing expected disagreement is then defined as $sED(P, Q) = \frac{1}{n} \sum\limits_{i=1}^{n} (1 - A_i(P_i Q_i))$.

##### S2.3.2. Evaluation metrics for minimum spatial migration

We begin with the weighted squared-change parsimony cost on continuous characters defined in [Maddison, 1991]. Given $(x, y)$ coordinates for each cell's centroid [Chadly et al., 2024], we compute the following independently for each coordinate,

$$\min_{\ell:V_T\to\mathbb{R}^d} \sum_{(u,v)\in E_T} w_v \left\|\ell(u)-\ell(v)\right\|^2, w_v = \frac{1}{\delta_e}$$

and where $\delta_e$ is the branch length on edge $e$.

As discussed in [Maddison, 1991], the optimal ancestral labeling under the squared-change parsimony is also the maximum likelihood ancestral labeling under a model of Brownian motion

$$\sum_{e\in E_T} \log\left(\frac{1}{\sqrt{2\pi v_e}}\right) - \frac{1}{2}\min_{\ell:V_T\to\mathbb{R}^d} \sum_{e\in E_T} \frac{(x_e-y_e)^2}{v_e}$$

To compare the maximum likelihood on two different tree topologies, we normalize by the degrees of freedom (here defined by the number of internal nodes for which we solve for spatial annotations).

#### S2.4. PEtracer data

Measurements are color-scaled and summarized across channels into a single pixel-intensity value per hybridization probe. Each site can take one of 9 states, and each observation is summarized into a pixel intensity vector $x \in \mathbb{R}^9 \cup \{?\}$, where "?" denotes that no fluorescence was observed at the site. As each of the three target sites takes different edit states, and following the original PEtracer discriminative classifier [Koblan et al., 2025], we produce 27 density estimates in total.

Note that the PEtracer tree uses weighted Hamming distance (wHD) to compute a distance matrix, then runs Neighbor-Joining to produce an unrooted tree topology. The tree is rooted using the default heuristic in Cassiopeia [Jones et al., 2020] and branch lengths in time units are estimated using ConvexML Prillo et al. [2025]. The tree is saved within the 'h5td' as a 'networkx' digraph, and was extracted with the 'time' unit branch lengths for our analyses.

We focused on three experiments from the PEtracer lineage tracing system. We refer to the first experiment as PEtracer-sequencing, as it has both sequencing and imaging-based readout of preedited cells. To the best of our knowledge, the PEtracer discriminative classifier was trained on this dataset, using either sequencing readout or the preedited states as the true labels. Thus, given the training data with paired imaged and true genotypes, the PEtracer discriminative classifier was trained to predict genotype calls from the imaged observations. We detail the density estimation process first, before discussing the analysis of the additional datasets.

##### S2.4.1. PEtracer density estimation

For the purposes of evaluating LAML-Pro on this first PEtracer-sequencing dataset with paired sequencing and imaging observations, we split the dataset 70/30 for training and test, leaving 4,724 cells for training and 2,025 cells for evaluation. In contrast, the original PEtracer paper used the whole PEtracer-sequencing dataset on 6,749 cells and 96 recordable sites to train a classifier mapping imaged observations to the true edited state. Thus, we split into training/test only for this first evaluation on PEtracer-sequencing. For subsequent evaluations on PEtracer-colonies and PEtracer-barcodes, we followed the PEtracer pipeline in leveraging all of the paired sequencing and imaging data to train our densities.

The PEtracer data was not well fit by standard parametric distributions (Supplementary Figure S8; we show only a single site for brevity) but provide the other plots on the reproducibility repository. We perform z-normalization on the published training data and call this data matrix $X$. We learn non-parametric kernel densities using statsmodels with a Gaussian kernel, assuming anisotropic, diagonal covariance, and solving for optimal bandwidth using the default (Scott's normal reference rule per dimension): $h_j = 1.06 \cdot \text{std}(X_j) \cdot n^{-1/(4+d)}, \forall j \in \{1, \ldots, d\}$, where $X_j$ is the data vector corresponding to the $j$th feature. Thus in this implementation, **bandwidth is scaled by the variance in each dimension**; please note that this is not equivalent to the default implementations in scipy and scikitlearn. We further note that this approach to density estimation is **not rotation invariant**.

We confirm that the intrinsic dimensionality of the data is 9 dimensions (Supplementary Figure S9). This is consistent with our mechanistic understanding, since the 9 pixel features correspond to the 9 hybridization probes matching the 9 possible states.

We also generated these scree plots after training the KDEs on the full paired sequencing/imaging dataset; these looked the same and so we have omitted them for brevity.

##### S2.4.2. PEtracer: sequencing validation

We use the PEtracer-sequencing dataset to compare the genotypes inferred by LAML-Pro, by sequencing, by PEtracer, and by taking the genotype corresponding to the brightest fluorescence measurement (reasonable baseline also used in Koblan et al. [2025]). We observe LAML-Pro genotypes to look the most similar to the sequencing dataset (Supplementary Figure S11); notably, LAML-Pro removes spurious genotype errors visible in the PEtracer approach and baseline approach of taking the genotype corresponding to the brightest measurement.

Next we evaluated whether LAML-Pro could impute genotypes at sites missing both imaged and sequenced observations. There were twelve such sites (Supplementary Figure S10). We hypothesize LAML-Pro did not impute more genotypes due to the combination of consistently high rates of missing at each site as well as the model misspecification – all sites are completely phylogenetically uninformative, so that the shared missing data is preserved as phylogenetic signal.

##### S2.4.3. PEtracer: colonies and barcode experiments

For the following two experiments, we trained the densities on all available labeled data from the above paired sequencing/imaging data. We observed similar quantile-quantile and scree statistics.

**PEtracer: colonies experiment** We refer to this dataset as PEtracer-colonies. In the PEtracer-colonies experiment, we are given imaged observations for 832 cells over 48 sites. The PEtracer analysis decided a $> 60\%$ cutoff for cassette detection (meaning that more than 60% of target sites must be present), throwing out cells which fell below this detection rate. Since the choice of a threshold cutoff is somewhat arbitrary, we explored LAML-Pro's ability to handle additional genotype uncertainty and missing data by relaxing this threshold to 50%.

We evaluated the genotype differences between LAML-Pro and the PEtracer approach (Supplementary Figure S12), and found that LAML-Pro genotype frequencies are similar to PEtracer genotype frequencies, and that LAML-Pro genotype frequences at sites with no observations were similar to LAML-Pro genotype frequencies at sites with observations (Supplementary Table S2).

We also ran our reimplementation of LAML on this dataset, by passing the PEtracer genotypes as input to LAML. Since LAML does not fix genotype errors, we do not include LAML for the genotype evaluations. We include LAML's statistics on spatial concordance in the main text.

**PEtracer: barcode experiment** We refer to this dataset as PEtracer-barcodes. In the PEtracer-barcodes analysis, cells were tagged with heterogeneous lentiviral barcode integrations at two different stages of growth. Then, the PEtracer paper evaluated the quality of the tree reconstructed with the PEtracer via consistency with the orthogonal measurement of lentiviral barcode labels in each cell. We focused on Clone 4, with 3,108 4T1 cells which was also analyzed in detail in the PEtracer paper Koblan et al. [2025]. LAML-Pro was given 25,000 NNI iterations to improve the published PEtracer tree Koblan et al. [2025], and finished in 18 hours. Although this experiment had sequencing-based readout, only the character matrix (with genotypes already called) was published. We made our best attempt to reprocess the sequencing data from the provided GEO accession references following Koblan et al. [2025], but the published pipeline required a custom reference which was not included. Thus, we processed the character matrix into an observation matrix, using a range of "error probabilities"; that is – $\eta = [0.01, 0.03, 0.05, 0.07, 0.10]$, and we took the tree with the highest likelihood. Note that this error probability is spread uniformly across all other genotypes possible at each site.

In this experiment, cells carry *static barcodes* that were irreversibly incorporated at two known timepoints. The static barcodes are designed so that cells with the same static barcode likely descend from a single ancestral integration event, so that reconstructed trees should group cells sharing the static barcodes together.

More specifically, the cell population is infected with a large library of lentiviral barcodes at two timepoints (at days 5 and 7, called "puro" and "blast" respectively) Koblan et al. [2025]. Thus, due to the size of the lentiviral barcode library, it is unlikely that two cells sharing a lentiviral barcode come from two separate infections with the same barcode. We note that these annotations are imperfect, as not every cell is observed with a lentiviral barcode at both timepoints.

To evaluate the LAML-Pro tree, we report the Wagner parsimony score, computed on each timepoint separately. The theoretical minimum is computed as the number of unique lentiviral barcodes at a given timepoint (subtracting one for minimum parsimony score, and adding one for the missing label state).

LAML-Pro produced a tree with plausibly fewer integration events of the static barcodes compared to the PEtracer tree, taking an average of 2.6 seconds per NNI iteration. At Timepoint 1, LAML-Pro required only 42 events compared to 53 events for PEtracer (theoretical minimum = 29 events), and at Timepoint 2 LAML-Pro required 195 events compared to 236 events for PEtracer (theoretical minimum = 96 events).

#### S2.5. baseMEMOIR data

Each of the 396 dinucleotide target sites is in one of five possible states: the 'AA' unedited state, the three edited states ('AG', 'GG', 'GA'), or the silenced state. Mathematically, these states are represented by a hidden alphabet $\mathcal{A}_H = \{0, 1, 2, 3, -1\}$, where "0" corresponds to the unedited state and "-1" corresponds to the silenced state. Pixel intensity features are summarized into a pixel intensity vector $\mathbf{x} \in \mathbb{R}^{16}$ and the observed alphabet is represented as $\mathcal{A}_O = \mathbb{R}^{16} \cup \{?\}$, where "?" denotes that no fluorescence was observed at the target site. Following Chadly et al. [2024], the measured intensities in each pseudocolor are summarized into four features (sum pixel intensity, median pixel intensity, top 10% pixel intensity, and pixel intensity variance), so that each observation $\mathbf{x} \in \mathbb{R}^{16} \cup \{?\}$.

Using the manually labeled training data Chadly et al. [2024], we fit four state-specific conditional densities $f_{\mathbf{z}}(\mathbf{x})$, $\mathbf{z} \in \mathcal{A}_H \setminus \{-1\}$, that provide the density of observing the pixel intensity vector $\mathbf{x} \in \mathbb{R}^{16}$ conditioned upon the hidden state $\mathbf{z}$. To fit the conditional densities $f_{\mathbf{z}}$, we perform dimensionality reduction with PCA (see e.g. Silverman [2018]) followed by non-parametric density estimation using a Gaussian kernel using the statsmodels package Seabold et al. [2010] (see Supplementary Section S2.5.1).

For validation and model training, a subset of these target site observations were manually labeled with states in Chadly et al. [2024] and used to train a logistic classifier.

Although log-normal distributions have previously been used to model fluorescence intensities, the pixel intensity vectors $\mathbf{x} \in \mathbb{R}^{16}$ were poorly fit by parametric distributions ($\leq 10^{-7}$ Shapiro-Wilks p-values over all features after log transformation and Anderson-Darling values ranging from 1.0 to 324; see Supplementary Figure S14). Furthermore, distinct features, e.g. sum pixel intensity and pixel intensity variance, followed different distributions, hindering the parametric approach.

The four largest colonies are Colonies 7/8, 9/10, 2 and 5. However, we were only able to recover feature vectors for 38 cells in Colony 7/8 and 36 cells in Colony 9/10. Furthermore, after resolving cells which were observed twice (e.g. once in colony 7 and once in colony 8 but with distinct labels), we have similar numbers of cells as Colonies 2 and 5, which did not require us to resolve duplicates. Thus, we use Colonies 2 and 5 for further analysis to avoid confusion.

##### S2.5.1. baseMEMOIR density estimation

The baseMEMOIR data was not well fit by standard parametric distributions (Supplementary Figure S14, showing only for the unedited state for brevity). We provide similar quantile-quantile plots for all states in the reproducibility repository. This fits expectations, since the 16 features include 4 summary features for each state (e.g. sum pixel intensity, variance in pixel intensity, median pixel intensity, top 10% brightest pixel intensity). We are given the following numbers of training samples; AA: 3658, AG: 2125, GG: 2067, GA: 1748. We perform z-normalization on the $\log(1 + x)$ transformed data provided in the baseMEMOIR paper Chadly et al. [2024], and call this data matrix $X$. We learn non-parametric kernel densities using statsmodels with a Gaussian kernel, as was done for PEtracer (Supplementary Section S2.4.1).

One standard approach to identify the intrinsic dimensionality of the data is to perform dimensionality reduction and evaluate using a metric which is dimension-invariant. On the baseMEMOIR data, we have 16 features, which correspond to 4 summary measurements for each of 4 states. We additionally have a mechanistic reason to believe our data lies on a lower dimension, since the hybridization probes produce four "pseudocolors" corresponding to the four observable states. Additionally, the scree plot further illustrates that 4 principal components explain most ($\approx 94\%$ of the variance in the training data; see Supplementary Figure S15 (a)).

Recall that principal components analysis (PCA) transforms a data matrix $X \in \mathbb{R}^{(n \times 16}$ to the same target dimension 16 via an orthogonal transformation $V \in \mathbb{R}^{16\times 16}$. In this case, PCA amounts to a rotation of the data $Z = XV$. We first compare the density $f_{\text{og}}(x)$ estimated on the original data to the density $f_{\text{rot}}(z)$ estimated on the rotated data.

Standard approaches to evaluate densities in the same dimension involve variants of the integrated mean squared error or IMSE (more commonly referred to as the MISE) Seabold et al. [2010], Li et al. [2007]. For each state, we report the approximation of the ISME implemented in *statsmodels*, defined below:

$$ISME = \int \hat{f}(x)^2 dx - 2E_X[\hat{f}(x)]$$

Comparing KDEs estimated on $Z$ versus $X$, we find that for each state, the rotated data yield lower error (see Table S3). Intuitively, this confirms that rotating data to concentrate variance across dimensions improves the accuracy of our density estimates.

Since IMSE is not dimension-invariant, we evaluated density estimates across dimensions using classification error. Given data matrix $X$, we construct representations $Z$ with varying numbers of principal components (corresponding to dimensions $d \in [2, 16]$). We perform stratified 5-fold cross-validation (stratified since it aims to keep the proportions of class labels the same in each fold) on each $Z$, repeating the procedure three times with different random partitions (with fixed random seed for reproducibility). For each fold and class, we estimate the densities $f^d_{\text{AA}}(z), f^d_{\text{AG}}(z), f^d_{\text{GA}}(z), f^d_{\text{GG}}(z)$ at each dimension $d$. Test samples are classified by evaluating their density under each class, and assigning the label corresponding to the largest density. The classification accuracy shows little variation between dimensions 5 and 16 (ranging from 95.1% to 95.2% respectively; see Figure S16). We selected dimension 5 for subsequent analysis on baseMEMOIR.

##### S2.5.2. baseMEMOIR tree estimation

The baseMEMOIR tree is inferred under a phylogeography model assuming Brownian motion migration; spatial and cell state data were combined into a single BEAST2 analysis, and 5 independent MCMC chains were run to verify convergence to the same posterior distribution Chadly et al. [2024]. Following the original paper Chadly et al. [2024], we used the posterior consensus tree published for Trial 4.

##### S2.5.3. baseMEMOIR genotype imputation

Across both colonies, we observe a slight bias in LAML-Pro towards imputing the unobserved state at sites lacking any observations compared to sites with observations. For instance in Colony 5 we observe similar genotype frequencies over all observed sites (unedited: 70.91%, GG: 14.38%, GA: 8.23%, AG: 6.48%) compared to over all imputed sites (unedited: 72.35%, GG: 13.02%, GA: 6.68%, AG: 5.49%). One example of this is the clade of cells from 1 to 7 in Colony 5 contains missing observations which LAML-Pro imputes according to the cells' shared ancestry (Figure S17).

##### S2.5.4. baseMEMOIR spatial analysis

Although the baseMEMOIR tree and LAML-Pro trees are both ultrametric, the baseMEMOIR tree was published without the branch length above the root. This root branch length indicates the time of shared ancestry between the true origin cell and the shared lowest common ancestor of all observed cells, and is present in the LAML-Pro tree. Without a similar branch in the baseMEMOIR tree, and without annotation of the true spatial origin of the original cell, we decided to trim the root branch off LAML-Pro for the following spatial analysis. Note that branches smaller than $1e-03$ have been collapsed.

We scale both the LAML-Pro tree and the baseMEMOIR tree to $\tau = 72$ experimental hours Chadly et al. [2024], and scale again by empirical variance defined as $\hat{\sigma^2} = \frac{\text{Var}(x)+\text{Var}(y)}{2\tau}$. Thus, in each tree $\delta_e$ represents the expected Gaussian variance (or pixel movement per hour). The empirical variance was $\text{Var}(X) = 106,940, \text{Var}(Y) = 208,056$ for Colony 2 and $\text{Var}(X) = 169,707, \text{Var}(Y) = 109,683$ for Colony 5.

The ratio of standard deviations $\sqrt{\frac{\mathrm{Var}(Y)}{\mathrm{Var}(X)}}$ yields 1.39 for Colony 2, indicating about 49% more spread in the Y coordinate and 0.80 for Colony 5, indicating about 20% more spread in the X coordinate.

#### S2.6. Topology search robustness

Although LAML-Pro does not guarantee a globally maximum likelihood tree, on baseMEMOIR Colony 5 all twelve starting trees were refined into nearly the same topology – pairwise RF indicated a maximum difference of one bipartition (Supp. Fig. S20B). This lends credence to the final solution for Colony 5. In contrast, on Colony 2 LAML-Pro converged to several topologically different solutions, with a pairwise minimum RF distance of 0.52 (14 different bipartitions) and maximum of 0.92 (25 different bipartitions) across twelve starting trees (Supp. Fig. S20A). On PEtracer, in each case, the twelve topologies were also improved into several topologically different solutions.

#### S3. Supplementary Figures

| | Parameter Effect | | | Parameters | | |
|---|---|---|---|---|---|---|
| **Model condition** | **Heritable missing % of total matrix** | **Dropout missing % of total matrix** | **Pairwise distribution separability ($\rho$)** | **Heritable rate ($\nu$)** | **Dropout rate ($\phi$)** | **Distribution standard deviation ($\sigma$)** |
| h0d0r04 | 0% | 0% | 0.044 | 0 | 0 | 0.20 |
| h0d0r25 | 0% | 0% | 0.249 | 0 | 0 | 0.30 |
| h0d0r46 | 0% | 0% | 0.458 | 0 | 0 | 0.40 |
| h0d0r61 | 0% | 0% | 0.607 | 0 | 0 | 0.50 |
| h0d0r04 | 18.75% | 18.75% | 0.044 | 0.15 | 0.1875 | 0.20 |
| h0d0r25 | 18.75% | 18.75% | 0.249 | 0.15 | 0.1875 | 0.30 |
| h0d0r46 | 18.75% | 18.75% | 0.458 | 0.15 | 0.1875 | 0.40 |
| h0d0r61 | 18.75% | 18.75% | 0.607 | 0.15 | 0.1875 | 0.50 |

Table S1: Simulated data under 8 model conditions, with varying missing data and varied Gaussian variance $\sigma^2$ to control the genotype error.

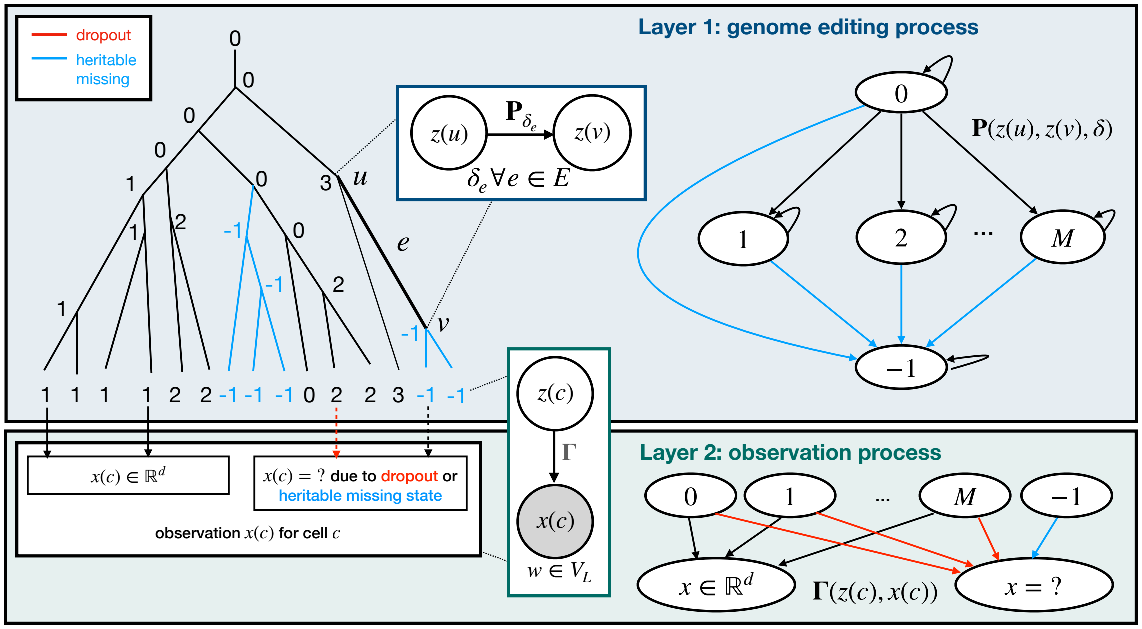

Figure S1: Model overview of PMMO. Layer 1 describes the genome editing process down a cell lineage tree, governed by transition probabilities $\mathbf{P}$ down each edge $e$. For each cell $u$, the hidden genotype is given by $z(u) \in \{0, 1, 2, \ldots, M, -1\}$, where $-1$ indicates the heritable missing state (transitions to this heritable missing state shown in blue). Layer 2 describes the observation process. The hidden genotype $z(u)$ produces observed measurement $x(c) \in \mathbb{R}^d$ with probability determined by the emission model $\mathbf{\Gamma}(z(c), x(c))$. The observations $\mathbf{x}_c$ take states in $\mathbb{R}^d \cup \{?\}$, where "?" indicates "missing observation." The transitions to the missing observation state correspond to stochastic dropout during observation (these transitions are shown with red edges).

| | ?/−1 | 0 | 1 | 2 | 3 | 4 | 5 | 6 | 7 | 8 | Total |
|---|---|---|---|---|---|---|---|---|---|---|---|
| PETracer | 0.2036 | 0.2788 | 0.0667 | 0.0499 | 0.0568 | 0.0506 | 0.0796 | 0.0257 | 0.0359 | 0.1525 | 1.0000 |
| LAML-Pro (obs) prop. over all sites | 0.0000 | 0.2774 | 0.0667 | 0.0497 | 0.0567 | 0.0507 | 0.0792 | 0.0250 | 0.0355 | 0.1554 | 0.7964 |
| LAML-Pro (obs) prop. over observed sites | nan | 0.3483 | 0.0838 | 0.0624 | 0.0712 | 0.0636 | 0.0995 | 0.0314 | 0.0446 | 0.1952 | 1.0000 |
| LAML-Pro (impute) prop. over missing sites | 0.0914 | 0.3404 | 0.0676 | 0.0534 | 0.0691 | 0.0464 | 0.1107 | 0.0206 | 0.0412 | 0.1592 | 1.0000 |

Table S2: PEtracer-colonies. Inferred genotype frequencies from PEtracer and LAML-Pro, across 9 possible states and the missing state. The missing state is given by '?' for PEtracer and −1 for LAML-Pro. The first two rows show a comparison between PEtracer and LAML-Pro genotype frequencies over all sites. The second two rows compare LAML-Pro genotype frequencies on all sites with observations, to LAML-Pro on all *imputed* genotype frequencies on sites where observations are missing.

| Label | ISME on KDE (Z) | ISME on KDE (X) |
|---|---|---|
| AA | -4.24389 | -0.00038 |
| GG | -2.67028 | -0.00008 |
| AG | -5.71509 | -0.00023 |
| GA | -0.81250 | -0.00004 |

Table S3: Density estimation error (Python package *statsmodels* definition of ISME) comparing density estimates on rotated data $Z$ to density estimates on original data $X$.

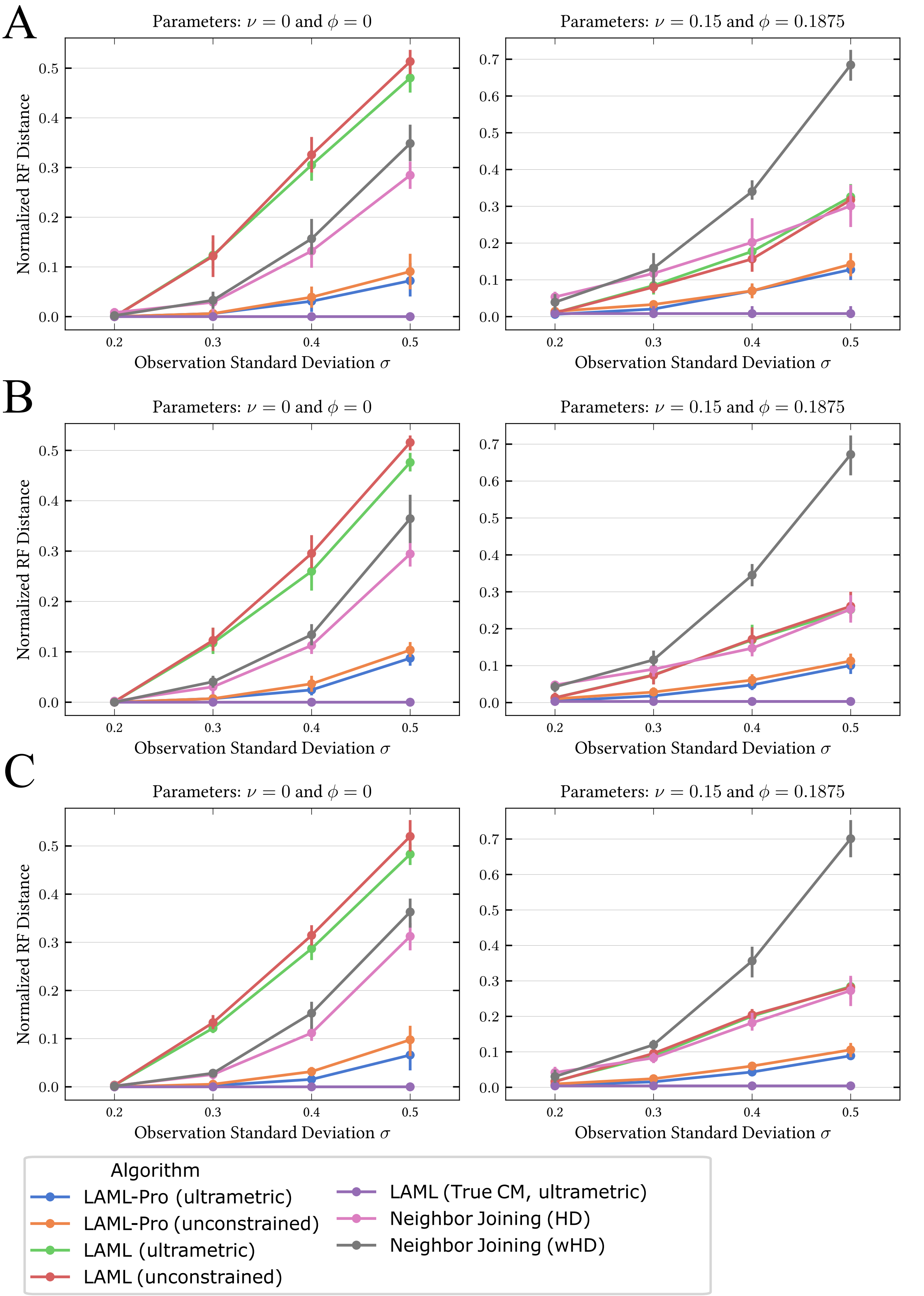

Figure S2: Evaluation of topology reconstruction accuracy on simulated datasets with $N = 100$ (A), $N = 200$ (B), and $N = 300$ (C) cells. For each dataset size, we compare LAML-Pro and several others. The left column shows the model condition with no missing data, and the right column shows the model condition with heritable missing rate $\nu = 0.15$ and dropout probability 0.1875. In all panels, the x-axis varies the observation standard deviation $\sigma$, and the y-axis reports the normalized RF distance between the inferred

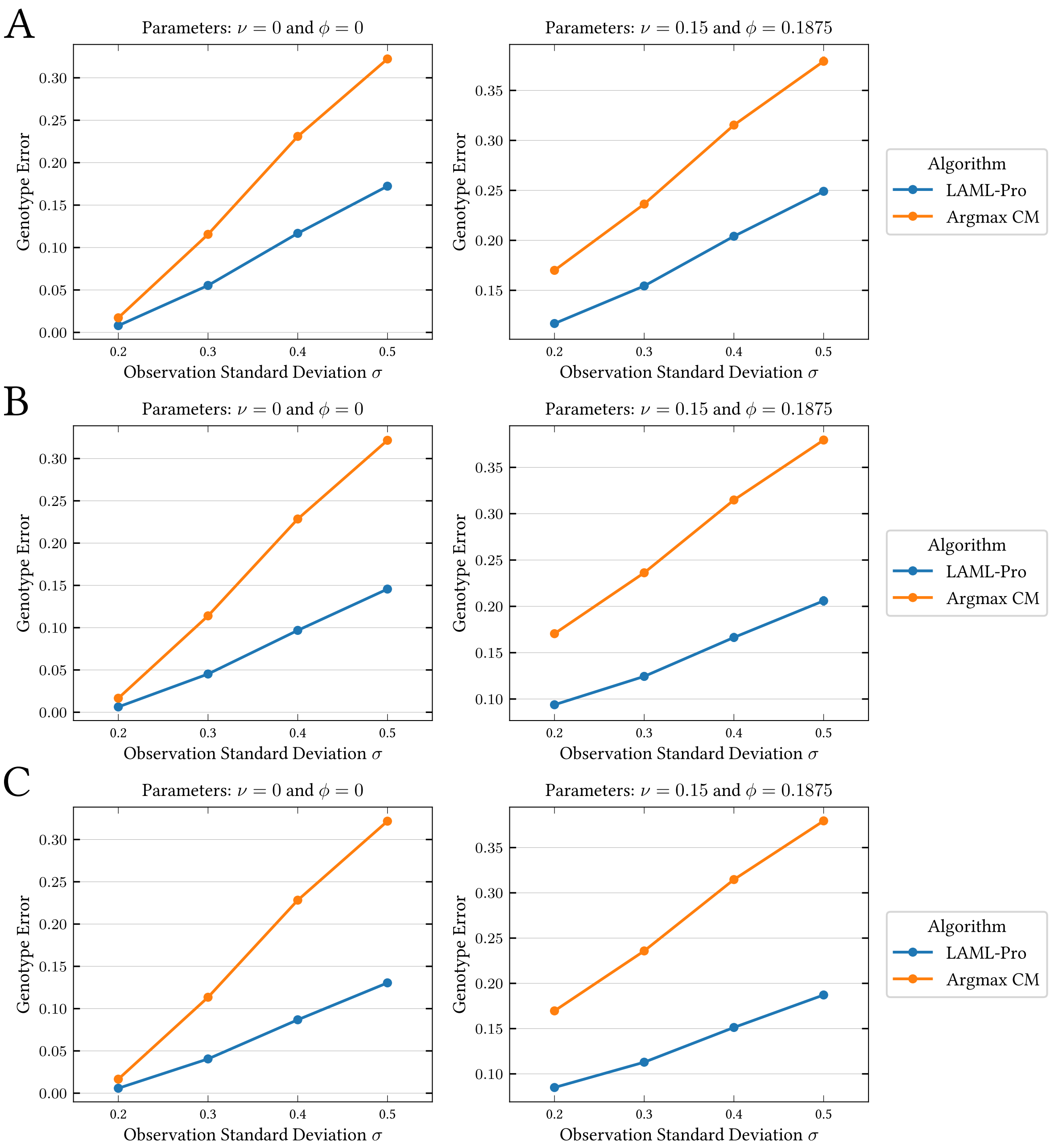

Figure S3: Evaluation of genotype error for $N = 100$ (A), $N = 200$ (B), and $N = 300$ (C) cells. For each dataset size, we compare LAML-Pro and several others. The left column shows the model condition with no missing data, and the right column shows the model condition with heritable missing rate $\nu = 0.15$ and dropout probability 0.1875. In all panels, the x-axis varies the observation standard deviation $\sigma$, and the y-axis reports the genotype error computed over all sites, including those without observations. Error bars indicate standard deviation over five independent replicates.

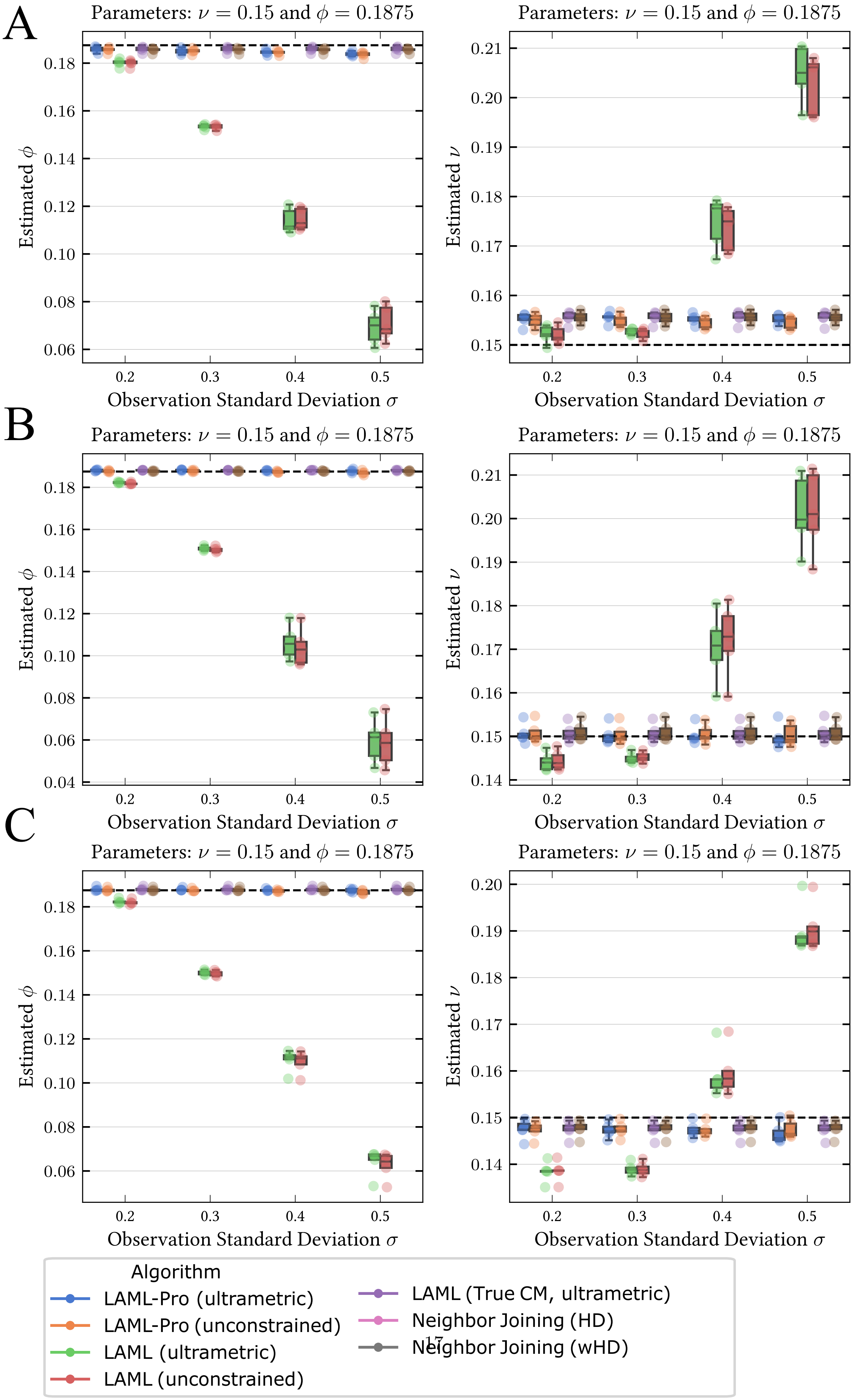

Figure S4: Evaluation of parameter estimation under the model condition with missing data (true values

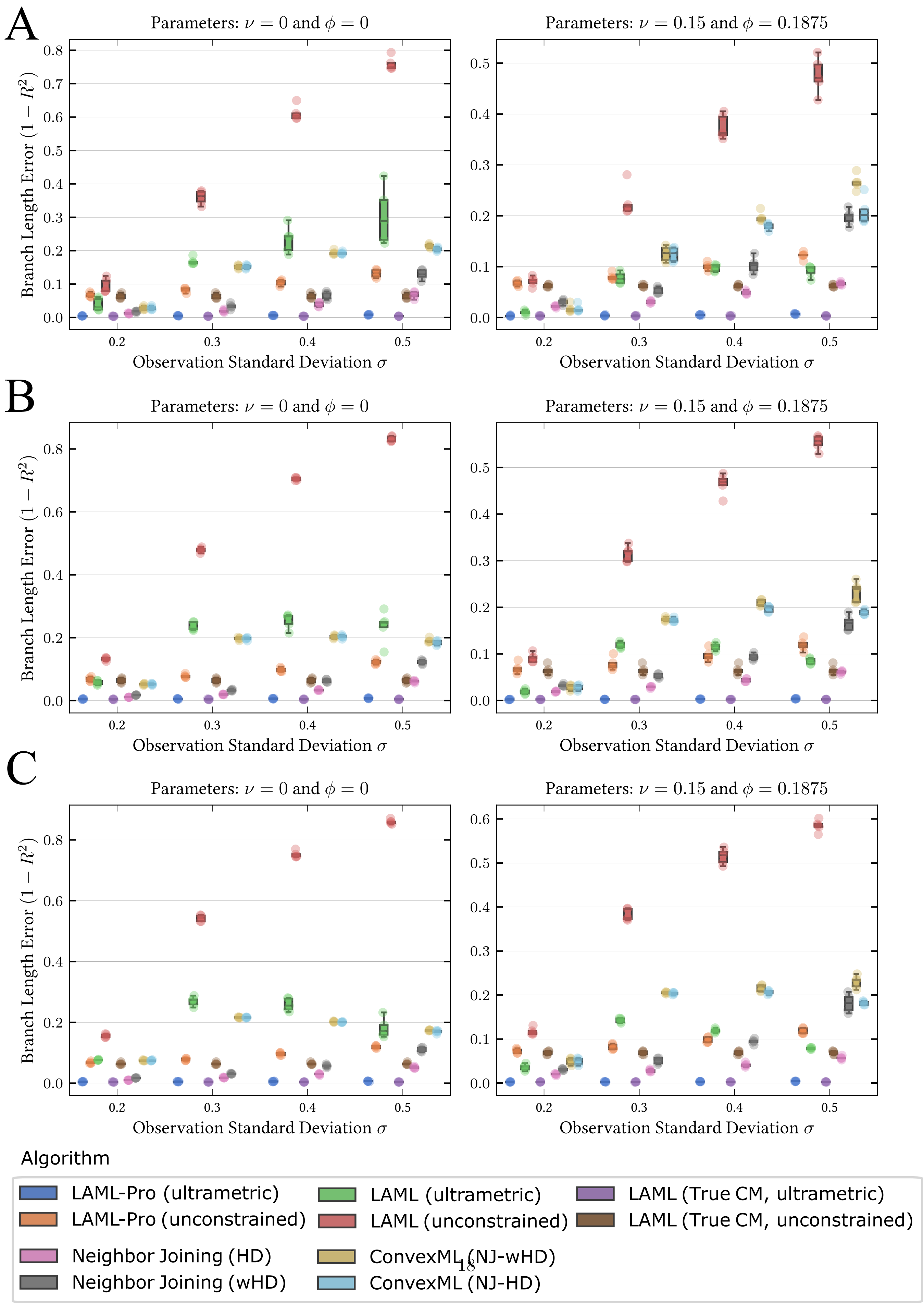

Figure S5: Evaluation of estimated branch lengths on simulated datasets with $N = 100$ (A), $N = 200$ (B), and $N = 300$ (C) cells. For each dataset size, we compare LAML-Pro and several others. The left

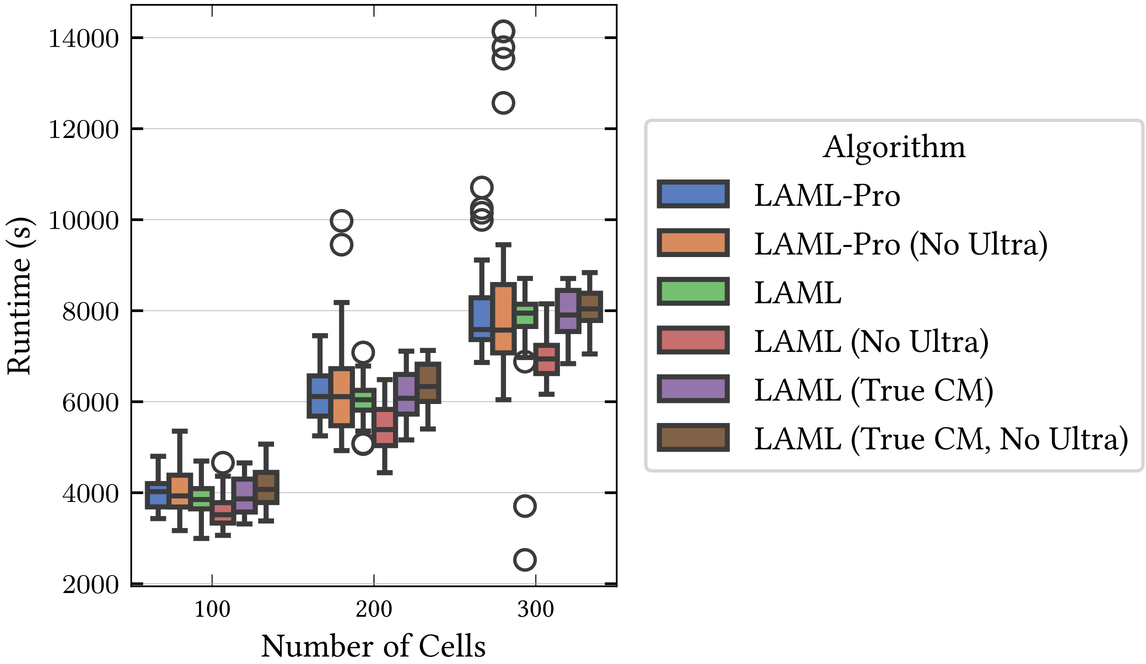

Figure S6: Topology search runtime evaluation on simulated datasets. The x-axis shows the number of cells, and the y-axis shows the runtime (seconds). We compare two ways of running LAML-Pro against two ways of running our re-implementation of LAML and LAML (True CM). All methods were given 5,000 NNI iterations to improve the starting trees.

| **Genotype Call Method** | **Colony** | **Missing/ Silenced** | **Unedited (AA)** | **GG** | **GA** | **AG** | **Row Total** |
|---|---|---|---|---|---|---|---|
| baseMEMOIR | 2 | 0.2995 | 0.4779 | 0.1238 | 0.0364 | 0.0624 | 1.0 |
| LAML-Pro (obs) over all sites | 2 | 0.0 | 0.5620 | 0.0689 | 0.0336 | 0.0360 | 0.7005 |
| LAML-Pro (impute) over all sites | 2 | 0.0202 | 0.2520 | 0.0143 | 0.0068 | 0.0061 | 0.2995 |
| LAML-Pro (obs) over observed sites | 2 | / | 0.8022 | 0.0984 | 0.0479 | 0.0514 | 1.0 |
| LAML-Pro (impute) over missing sites | 2 | 0.0675 | 0.8415 | 0.0478 | 0.0228 | 0.0205 | 1.0 |
| baseMEMOIR | 5 | 0.2435 | 0.4666 | 0.1540 | 0.0613 | 0.0745 | 1.0 |
| LAML-Pro (obs) over all sites | 5 | 0.0 | 0.5364 | 0.1088 | 0.0623 | 0.0490 | 0.7565 |
| LAML-Pro (impute) over all sites | 5 | 0.0060 | 0.1762 | 0.0317 | 0.0163 | 0.0134 | 0.2435 |
| LAML-Pro (obs) over observed sites | 5 | / | 0.7091 | 0.1438 | 0.0823 | 0.0648 | 1.0 |
| LAML-Pro (impute) over missing sites | 5 | 0.0246 | 0.7235 | 0.1302 | 0.0668 | 0.0549 | 1.0 |

Table S4: Comparison of genotype calls from LAML-Pro and baseMEMOIR across two colonies. LAML-Pro is split into two cases; genotype calls made at sites with observations (LAML-Pro (obs)) and sites without observations, where LAML-Pro imputes genotypes (LAML-Pro (impute)). The proportion of sites in each genotype state is reported over all sites. Missing data are either imputed or inferred to be in the silenced state by LAML-Pro, so the Missing/Silenced category does not apply to LAML-Pro (obs) over observed sites (denoted by '/'). Proportions in the Missing/Silenced category should be interpreted as Missing for baseMEMOIR and Silenced for LAML-Pro, as basEMEMOIR does not impute missing data.

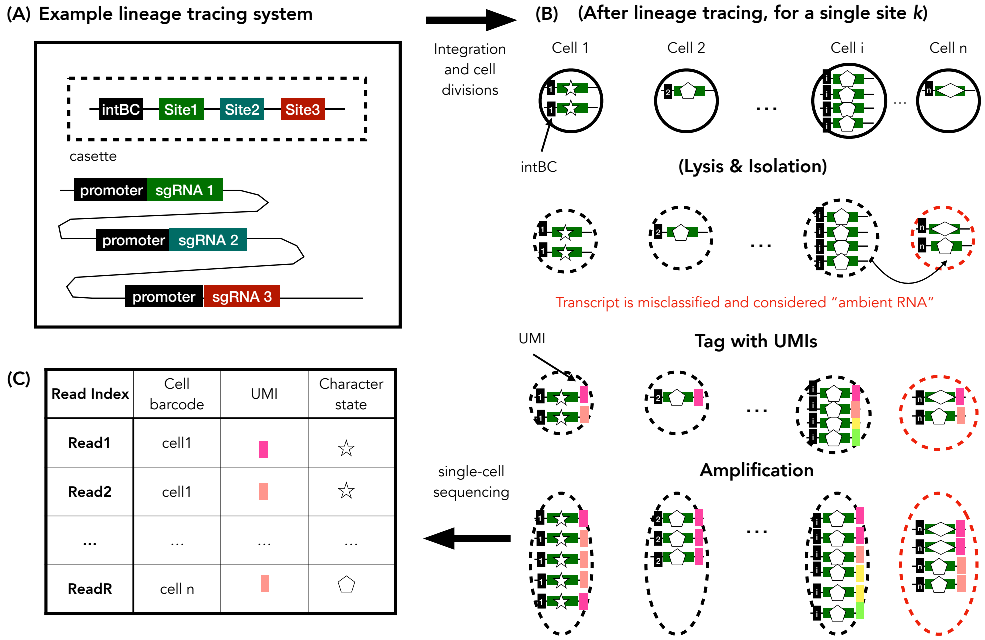

Figure S7: Illustration of "ambient RNA" as a source of sequencing error. (A) An example lineage tracing system Chan et al. [2019], Yang et al. [2022], Koblan et al. [2025], with a cassette containing three sites, and where each site has a corresponding promoter. This lineage tracing system is integrated into the cell, and over several cell divisions, each site may acquire edits. Each site's promoter transcribes its respective site. (B) After the end of the lineage tracing experiment, each cell is lysed and tagged with cell barcodes (Lysis & Isolation). Each site is identified by the cassette barcode (intBC), and all transcripts for this site are grouped by cell and barcode. During this period, cells with many transcripts may "shed," and a few transcripts may be incorrectly classified as belonging to a different cell. Following this, individual molecules may be tagged with unique molecular identifiers (UMIs) prior to amplification (generating many reads per UMI) so that the transcripts can be read out. (C) The final product is a list of reads. Each read is from a unique molecule, shows the genomic sequence, and is assigned to a cell (with the cell barcode).

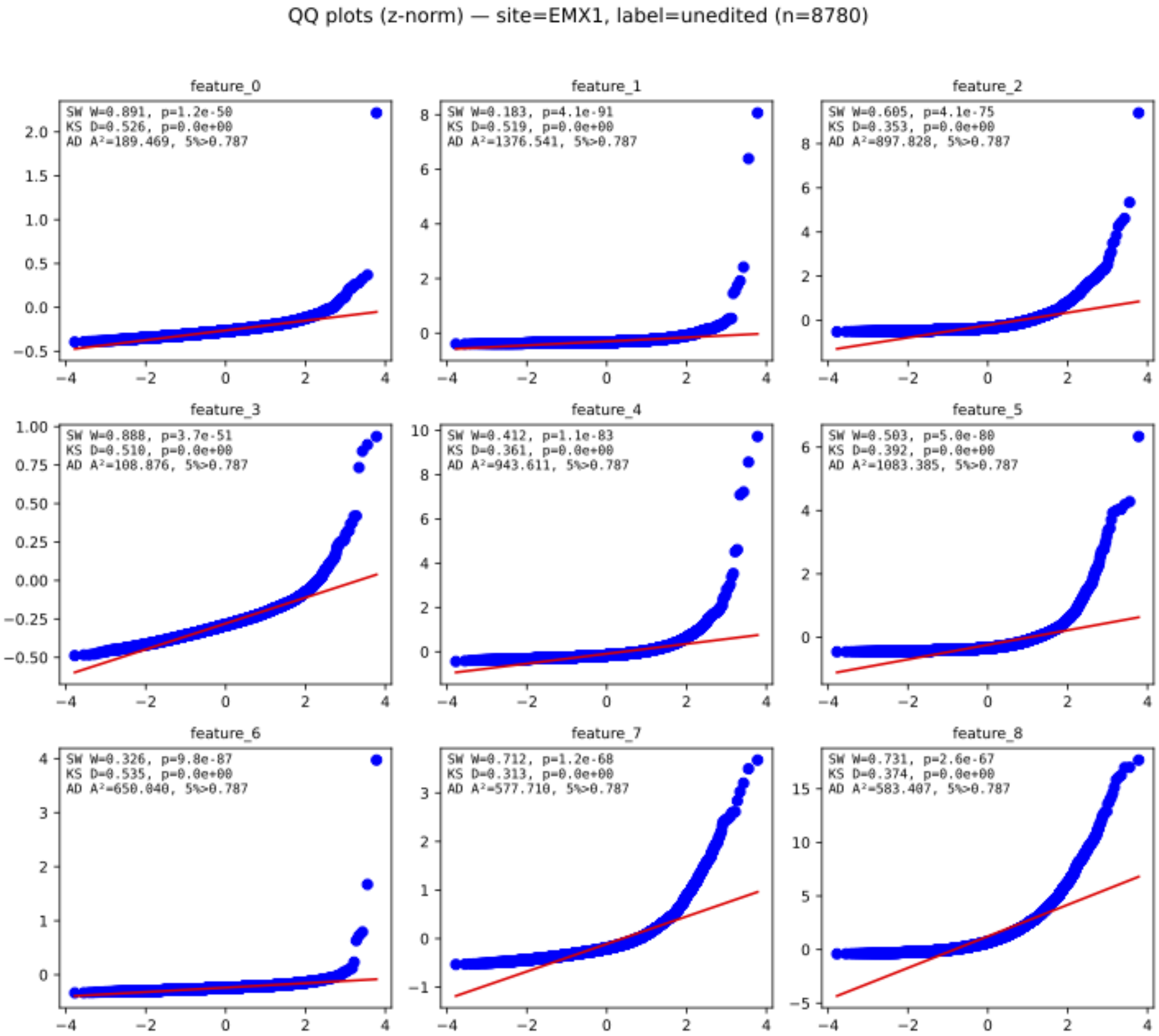

Figure S8: Quantile-quantile (QQ) plots on Z-normalized PEtracer observations at site EMX1 in the unedited genotype ($n = 8{,}780$) shown for each of the nine observed features. For each feature, we report three standard normality test statistics: Shapiro-Wilks (SW), Kolmogorov-Smirnov (KS), Anderson-Darling (AD).

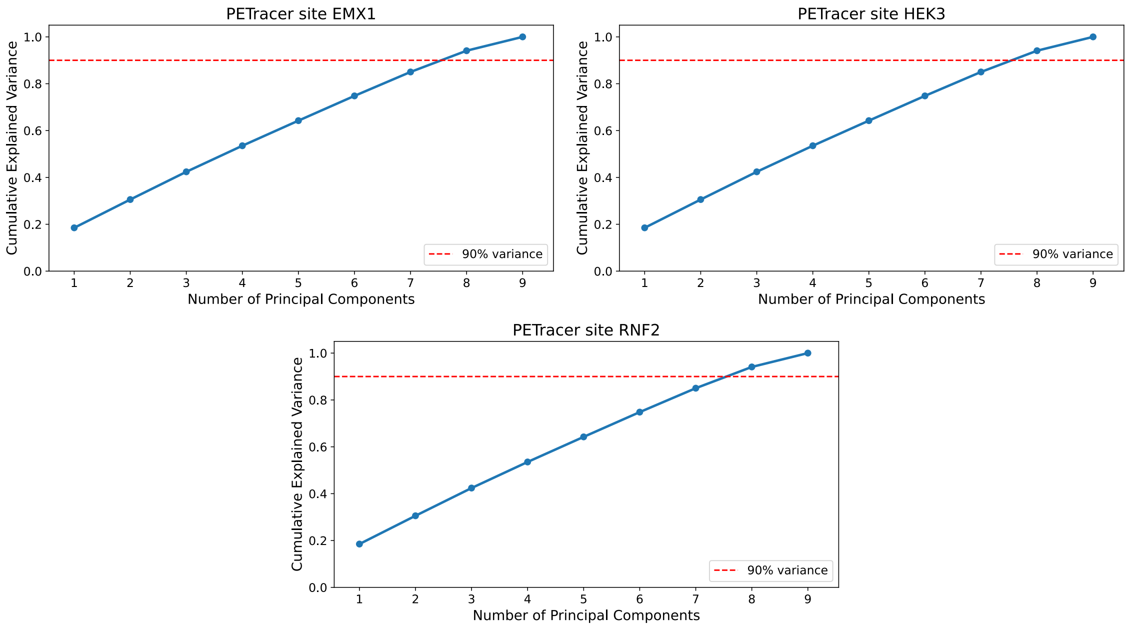

Figure S9: PEtracer-sequencing. Cumulative explained variance from principal component analysis (PCA) applied to the training subset (70/30 split) of PEtracer data for each of the three PEtracer target site designs: EMX1, HEK3, RNF2. As each site has a distinct edit alphabet, PCA was performed independently for each site. The red dashed horizontal line marks the 90% variance threshold.

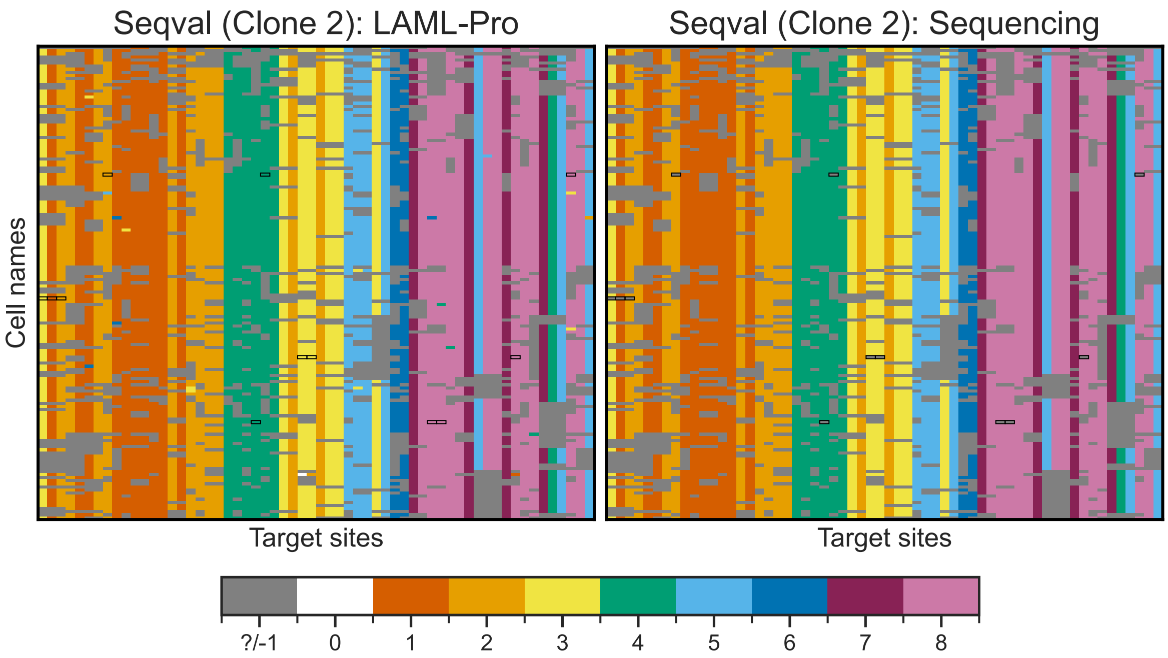

Figure S10: PEtracer-sequencing. Comparison of LAML-Pro estimated genotypes to the sequenced, predefined genotypes. Heatmaps show the inferred genotypes across all target sites for Clone 2, comparing LAML-Pro (left) with the predefined, sequencing-based genotypes (right). Black boxes indicate cells and sites at which LAML-Pro imputed genotypes where there were no sequencing or imaged observations. Note that −1 (gray) denotes missing observations (sequencing interpretation) and denotes the heritable silenced state for LAML-Pro. 0 is the unedited state (white). Edited genotypes (1–8) correspond to the site-specific edit alphabet.

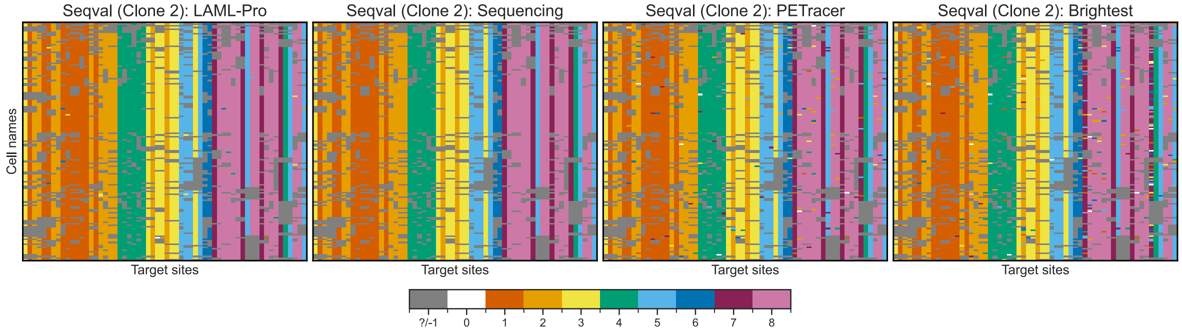

Figure S11: PEtracer-sequencing. Given PEtracer observations, we compare four approaches to genotype calls: LAML-Pro, sequencing calls as ground truth (filtered to be correct against the predefined edits), PEtracer, and a baseline of taking the state with the corresponding brightest fluorescence measurement. Note that −1 denotes missing observations (PEtracer, sequencing, brightest interpretation) and denotes the heritable silenced state for LAML-Pro. 0 is the unedited state.

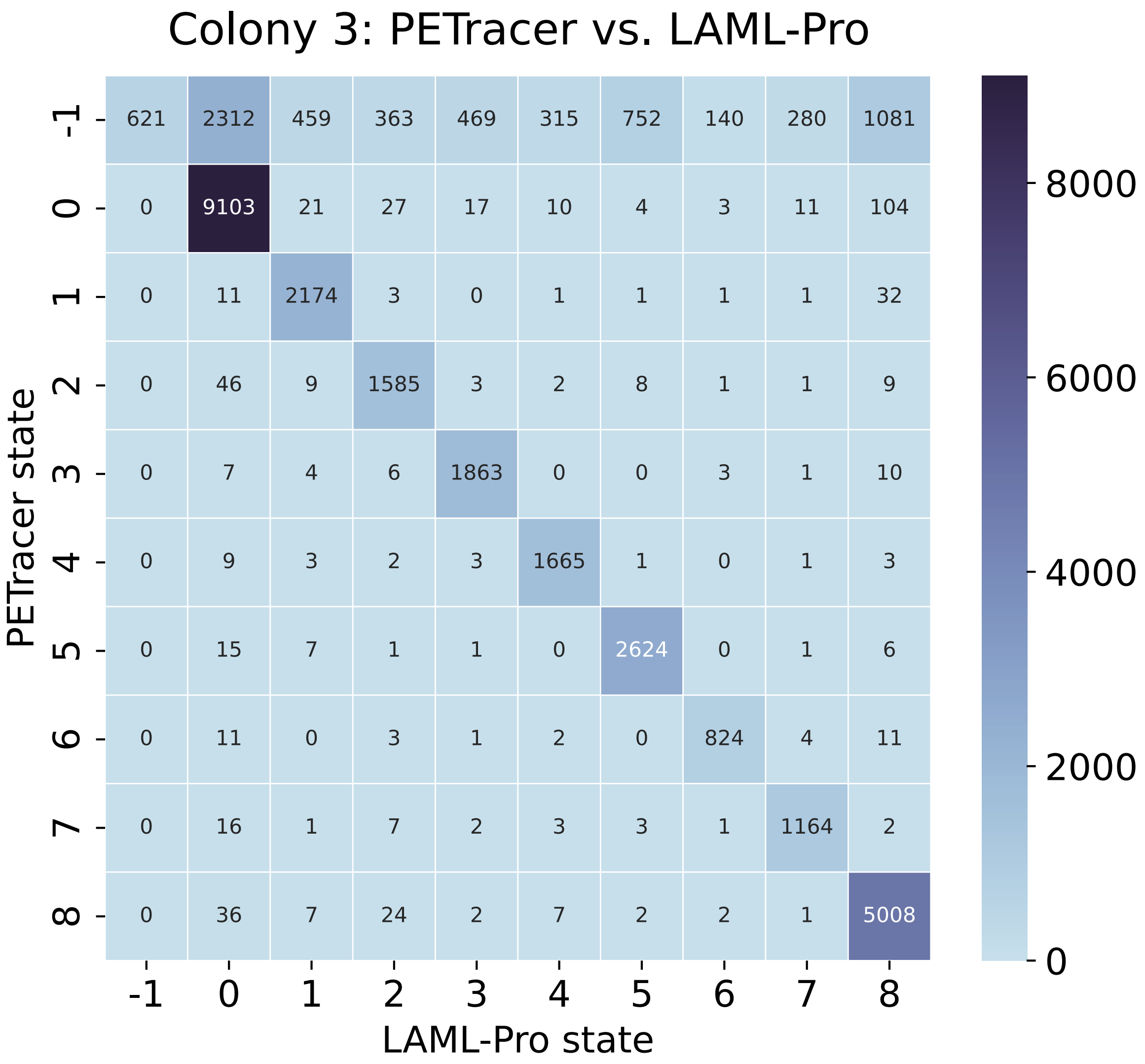

Figure S12: PEtracer-colonies. Confusion matrix comparing PEtracer genotype calls to LAML-Pro genotype calls for PEtracer Colony 3. Each cell shows the number of observations PEtracer assigned to a state (rows) and which LAML-Pro assigned to a state (columns). Note that −1 denotes missing observations along the rows (PEtracer interpretation), whereas −1 denotes the heritable silenced state along the columns (LAML-Pro interpretation). Additionally, 0 is the unedited state.

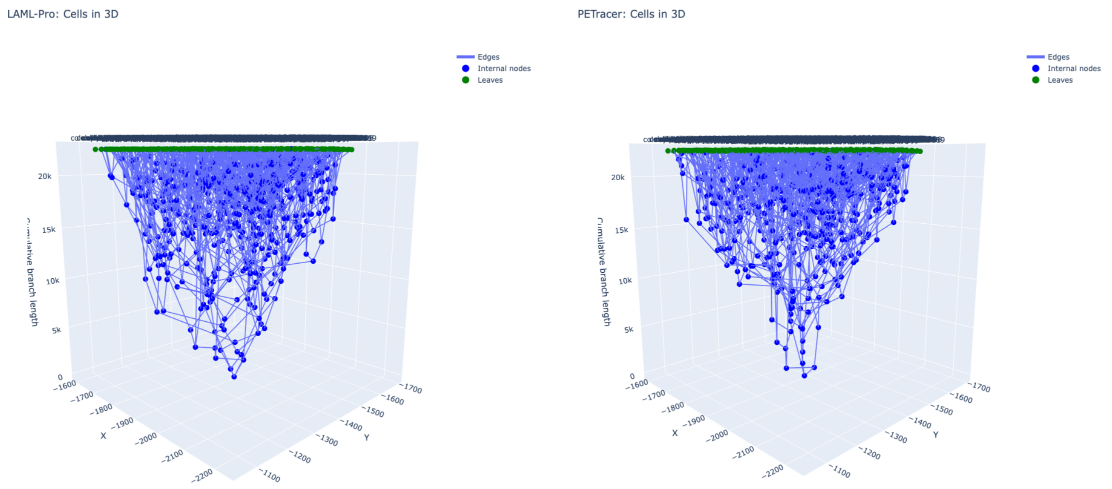

Figure S13: PEtracer Colony 3. The inferred LAML-Pro and published PEtracer trees visualized in 3D: $(x, y)$ are given by imaged spatial location and $z$ is given by the cumulative estimated branch length.

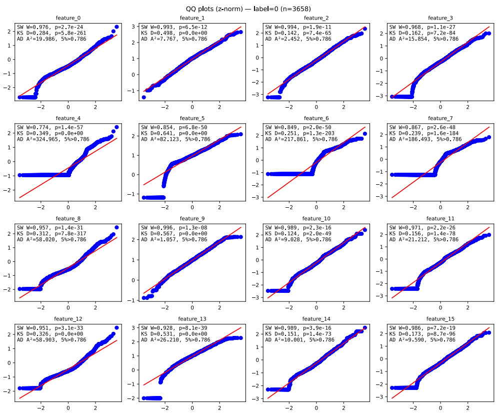

Figure S14: The training data provided by the baseMEMOIR authors was $\log(1+x)$ transformed. Typically, fluoresence intensities follow a log-normal distribution; here we provide the quantile-quantile plots comparing if our log-transformed feature data to the theoretical normal distribution.

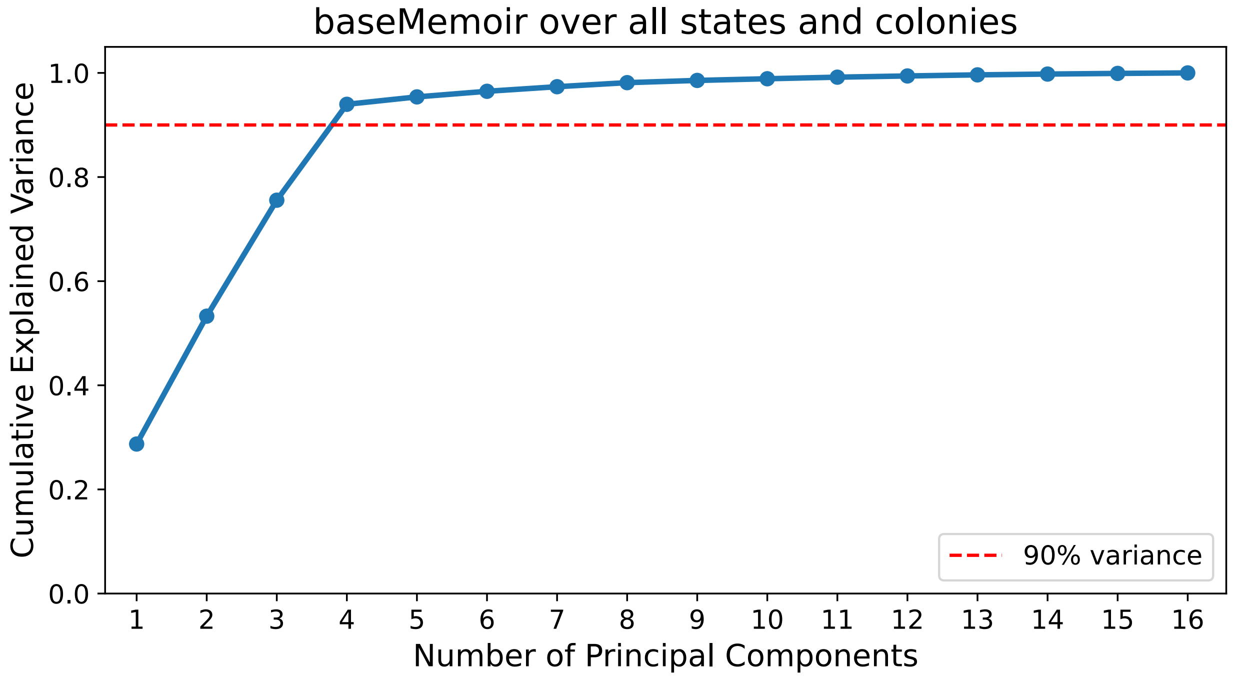

Figure S15: Evaluating dimensionality on the baseMEMOIR training data. Scree plot showing principal components on the x-axis and cumulative variance explained on the y-axis.

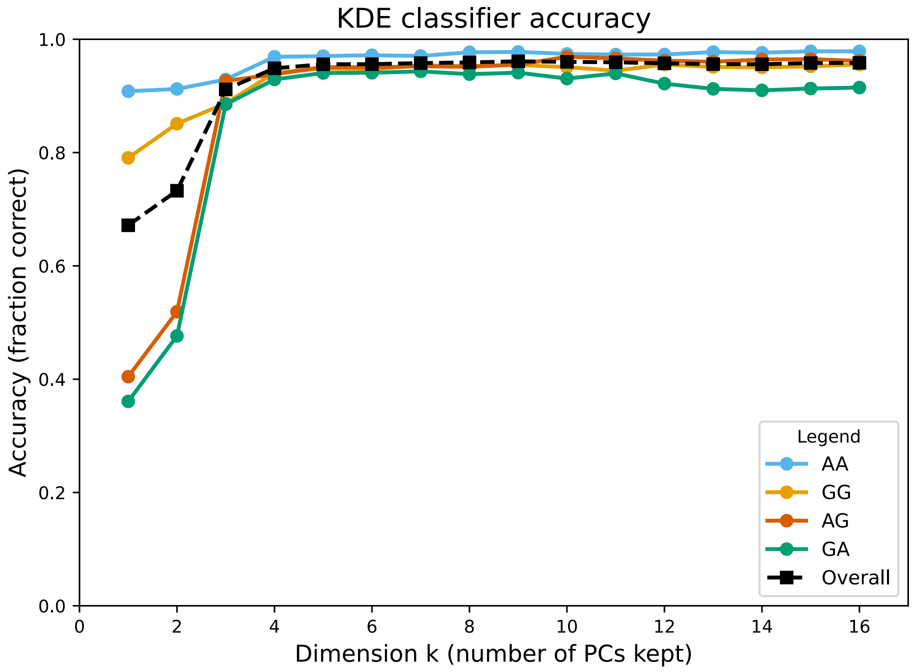

| Dimension k | Accuracy |
|---|---|
| 1 | 67.2% |
| 2 | 73.3% |
| 3 | 91.1% |
| 4 | 94.9% |
| 5 | 95.6% |
| 6 | 95.6% |
| 7 | 95.8% |
| 8 | 95.9% |
| 9 | 96.1% |
| 10 | 96.0% |
| 11 | 95.9% |
| 12 | 95.8% |
| 13 | 95.6% |
| 14 | 95.6% |
| 15 | 95.8% |
| 16 | 95.8% |

Figure S16: Classification performance on held out baseMEMOIR test data, evaluating kernel densities estimated on varying numbers of principal components.

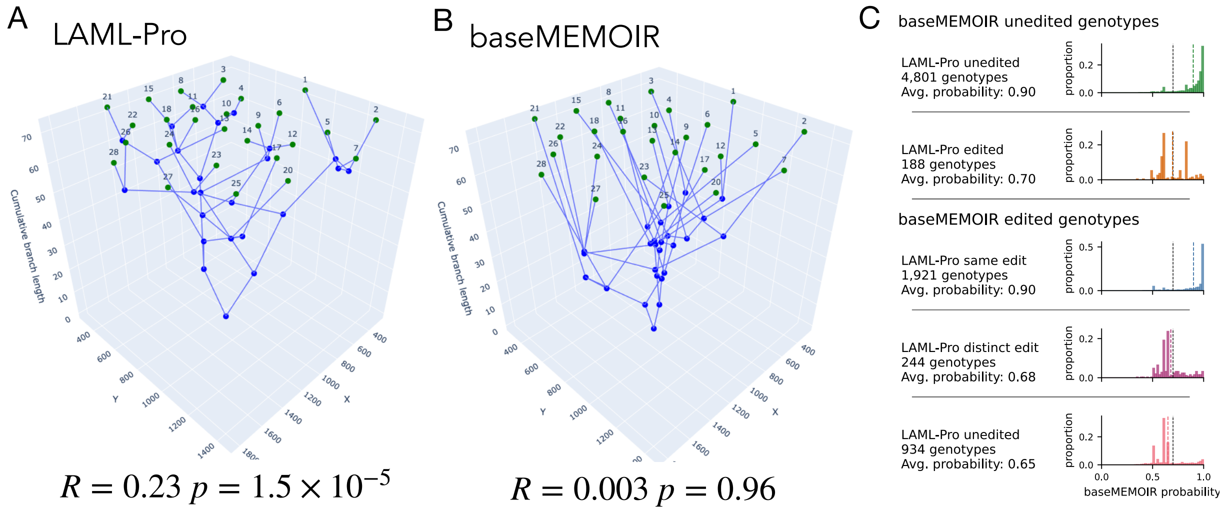

Figure S17: baseMEMOIR Colony 5. (A) The inferred LAML-Pro and published baseMEMOIR trees visualized in 3D (where $(x, y)$ are given by imaged spatial location and $z$ is given by the cumulative inferred branch length). (B) The baseMEMOIR and LAML-Pro genotypes on all sites with observations, broken into five categories. Each category is shown with a corresponding histogram illustrating the distribution of baseMEMOIR genotype probabilities.

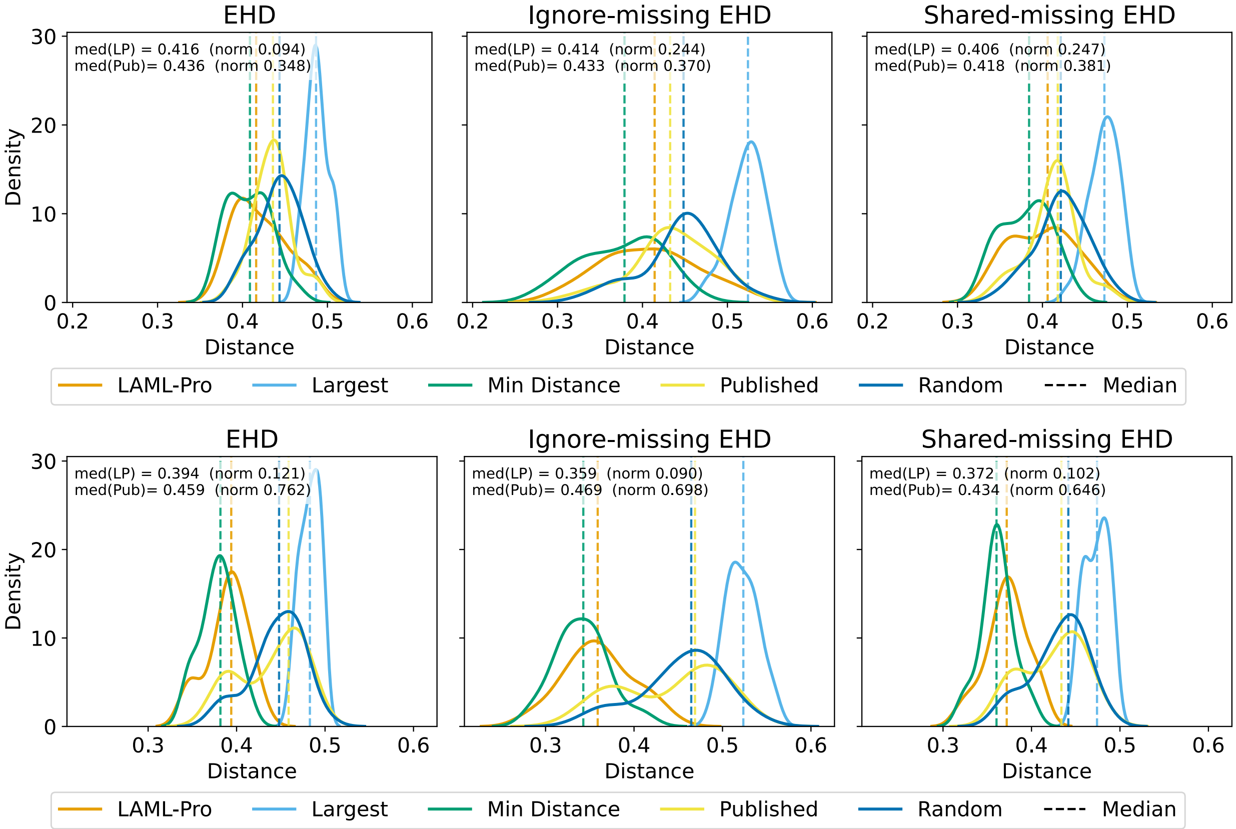

**A**

**B**

Figure S18: Evaluation of the LAML-Pro and baseMEMOIR trees in terms of concordance with the baseMEMOIR genotypes, on Colony 2 (A) and Colony 5 (B). We decompose each tree into sibling pairs, and evaluate the distribution of sibling pairs from each tree. Pairwise genotype distances computed by three different distance metrics: expected Hamming distance (EHD), ignore-missing EHD, shared-missing EHD). Solid curves show the distribution for several different definitions of cell pairs. Cell pairs are defined as follows: for each leaf, the LAML-Pro tree defines a closest sibling (orange), the published baseMEMOIR tree defines a closest sibling (yellow), the minimum distance cell genotype (green) and maximum distance cell genotype (light blue). We also sample random pairs as a second baseline (dark blue). Vertical dashed lines mark the medians for each corresponding method.

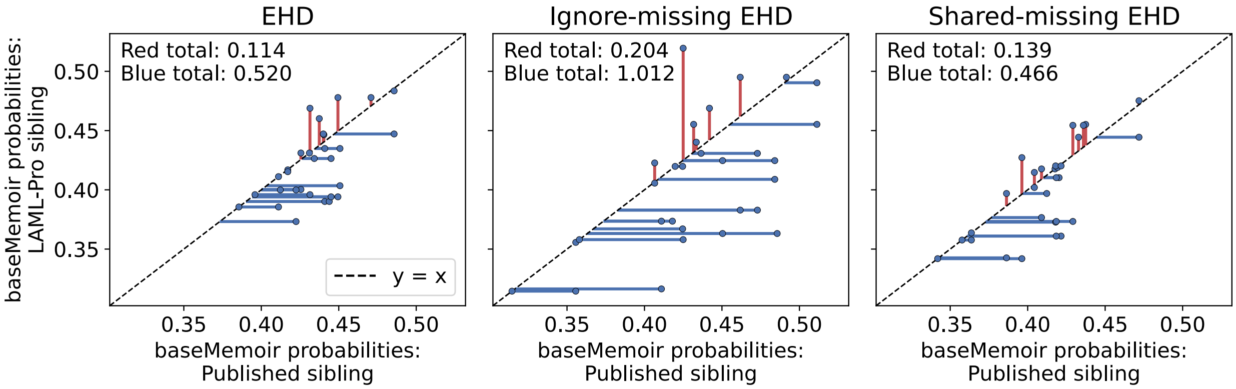

**A**

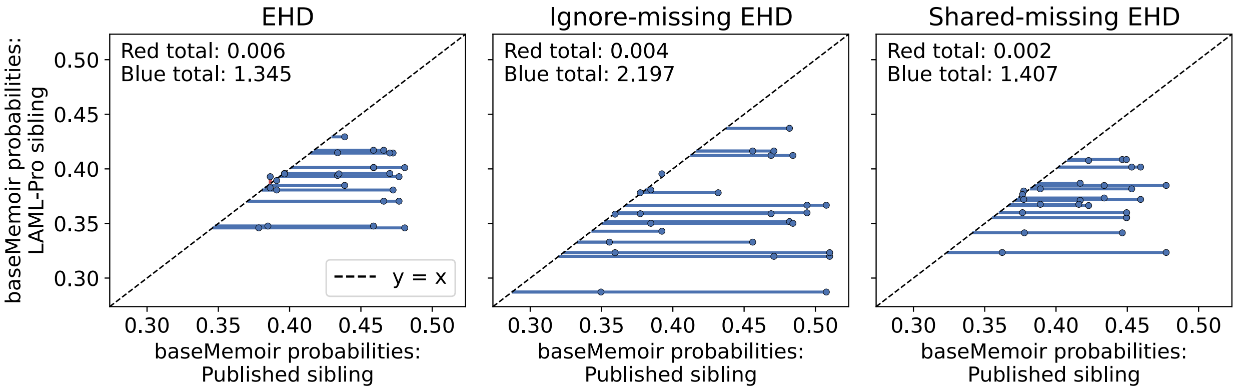

**B**

Figure S19: Scatterplot of the sibling pairs in the LAML-Pro and baseMEMOIR trees, evaluating in terms of concordance with the baseMEMOIR genotypes, on Colony 2 (A) and Colony 5 (B). For each leaf, we show the genotype distance to its closest sibling in the published baseMEMOIR tree (x-axis) and the closest sibling in the LAML-Pro tree (y-axis). We compute genotype distance using the baseMEMOIR probabilities, using three different distance metrics: expected Hamming distance (EHD), ignore-missing EHD, shared-missing EHD). The $x = y$ dashed line provides a baseline for each pair. Lines indicate genotype distance over the baseline incurred by LAML-Pro (red) and baseMEMOIR (blue) sibling pairs. Totals are reported in the top right corner.

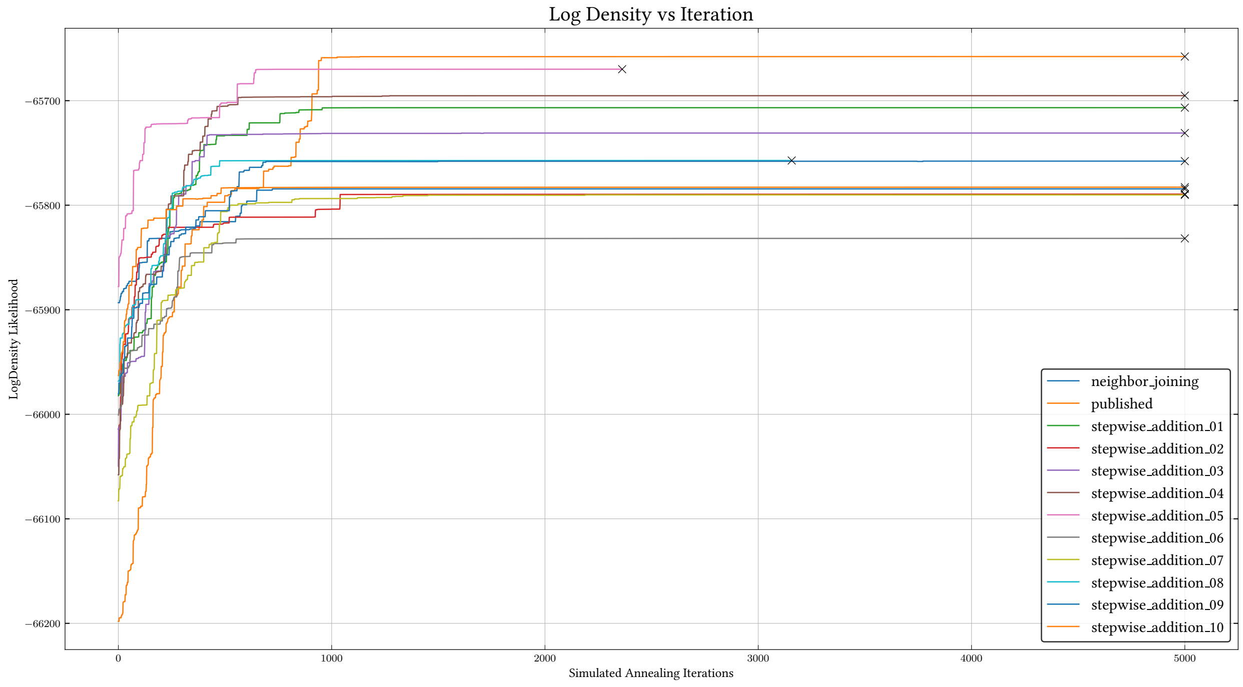

**A**

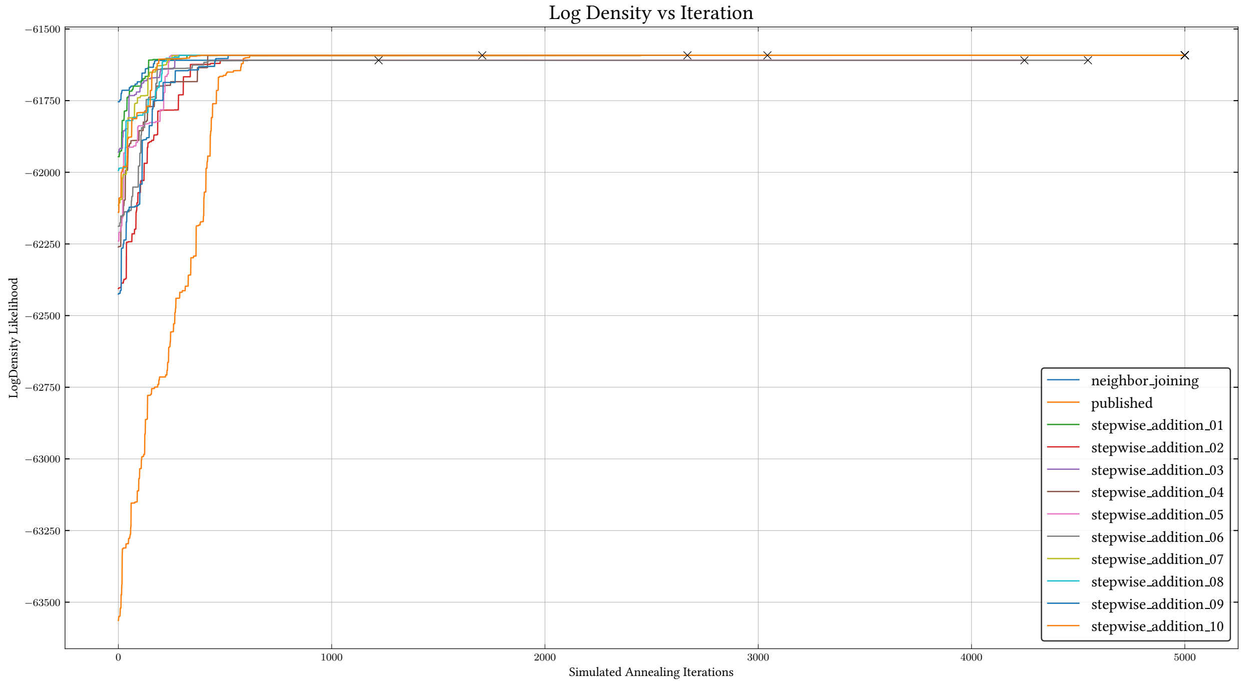

**B**

Figure S20: baseMEMOIR on Colony 2 (A) and Colony 5 (B). LAML-Pro log-density over 5,000 NNI iterations, given 11 initializations of the starting tree (10 parsimony-based initalizations and the published tree).
